# Accurate genotype-based demultiplexing of single cell RNA sequencing samples from non-human animals

**DOI:** 10.1101/2022.09.22.508993

**Authors:** Joseph F. Cardiello, Alberto Joven Araus, Sarantis Giatrellis, András Simon, Nicholas D. Leigh

## Abstract

Single cell sequencing technologies (scRNA-seq, scATAC-seq, etc.) have revolutionized the study of complex tissues and unique organisms, providing researchers with a much needed species agnostic tool to study biological processes at the cellular level. To date, scRNA-seq technologies are expensive, require sufficient cell quantities, and need biological replicates to avoid batch effects or artifactual results. Pooling cells from multiple individuals into a single scRNA-seq library can address these problems. However, sample labeling protocols for facilitating the computational separation of pooled scRNA-seq samples, termed demultiplexing, have undesirable limitations, particularly in resource-limited organisms. One promising solution developed for use in humans exploits the genetic diversity between individuals (i.e., single nucleotide polymorphisms (SNP)) to demultiplex pooled scRNA-seq samples. The use of SNP-based demultiplexing methods has not been validated for use in non-human species, but the widespread use of SNP-based demuxers would greatly facilitate research in commonly used, emerging, and more obscure species. In this study we applied SNP-based demultiplexing algorithms to pooled scRNA-seq datasets from numerous species and applied diverse ground truth confirmation assays to validate genetic demultiplexing results. SNP-based demultiplexers were found to accurately demultiplex pooled scRNA-seq data from species including zebrafish, African green monkey, *Xenopus laevis*, axolotl, *Pleurodeles waltl*, and *Notophthalmus viridescens*. Our results demonstrate that SNP-based demultiplexing of unlabeled, pooled scRNA-seq samples can be used with confidence in all of the species studied in this work. Further, we show that the only genomic resource required for this approach is the single-cell sequencing data and a *de novo* transcriptome. The incorporation of pooling and SNP-demultiplexing into scRNA-seq study designs will greatly increase the reproducibility and experimental options for studying species previously limited by technical uncertainties, computational hurdles, or limited cell quantities.

## Introduction

Over the last decade, single cell RNA sequencing (scRNA-seq) has exploded in popularity as a species agnostic tool for studying gene expression at the level of individual cells (Macosko et al. 2015; Klein et al. 2015; Han et al. 2018; Villani et al. 2017). Perhaps the biggest impact has been on species in which study at the cellular level was long difficult or impossible (i.e., all species other than rodents and primates). Though extremely powerful, the scRNA-seq approaches that have risen to prominence are expensive and low throughput for biological replicates. This is a serious drawback as the lack of biological replicates has also been shown to be a major cause of false discoveries (Squair et al. 2021; Hicks et al. 2018). To limit artifacts in scRNA-seq there is a critical need for approaches that allow for adequately powered experiments. Sample pooling is an effective means to increase biological replicate throughput while simultaneously decreasing batch effects and costs. Sample pooling methods can also enable scRNA-seq in the event of limited tissue or cell quantities and as an added benefit can facilitate doublet detection, a critical step when characterizing unique cell populations in complex tissues (DePasquale et al. 2019). Though using pooled scRNA-seq samples can provide these varied benefits, many of the methods available for demultiplexing pooled samples are only available and validated for samples with abundant cells and for well-studied species like humans or mice.

Methods for analyzing pooled data and for enabling the demultiplexing (also known as demuxing) of pooled scRNA-seq samples are varied in concept and accuracy and have been recently reviewed (Zhang et al. 2022). In developmental biology, which often requires small tissue samples, pooling of samples from multiple animals with no sample labeling method or intention of demultiplexing has become a standard practice. This method lacks advantages of true replicates because without demultiplexing the pooled samples there is no way to assess the data for representation of all samples, variation between samples, or cell clusters derived from one individual. Additionally, this approach lacks the ability to batch correct replicates and perform replicate strengthened differential expression analysis.

To enable these benefits, experimental protocols for demultiplexing of pooled scRNA-seq samples have been developed. These methods include pooling cells from transgenic animals or cell lines that express a distinct transgene (Lin et al. 2021) or oligonucleotide (Shin et al. 2019). The most popular and commercially supported method for scRNA-seq pooling is cell hashing, in which cells from each sample are labeled with antibodies (Stoeckius et al. 2018), lipids (McGinnis et al. 2019), or chemicals (Gehring et al. 2020) tethered to an oligonucleotide label that links gene expression data from each cell to the cellular origin (i.e., cell multiplexing oligonucleotide (CMO) label). A downside to these label-based approaches is that they each have varying degrees of cell assignment efficiency, require sufficient cell numbers, have costs associated with their application, are not compatible with all species, and have a chance of failing (Zhang et al. 2022).

In contrast to these experimentally-driven demultiplexing approaches, computational methods have been developed to demultiplex pooled human samples without any labeling regimen using the natural genetic differences between individuals. These approaches detect genetic differences between samples at sites of single nucleotide polymorphisms (SNPs), and implement demultiplexing based on differential distributions of these SNPs between samples. SNP-based approaches have been benchmarked and shown to be highly effective at separating human samples (Heaton et al. 2020; Y. Huang, McCarthy, and Stegle 2019; Kang et al. 2018; Xu et al. 2019; Weber et al. 2021). SNP-based demultiplexing has also been applied to demultiplex plasmodium samples, and across mouse strains (Mylka et al. 2022; Heaton et al. 2020). However, there are clearly limits to the utility of SNP-based demultiplexing, as it has been reported to not be possible to demux pooled samples from within the same mouse strain (Mylka et al. 2022; Heaton et al. 2020). Therefore it was unclear whether the success and accuracy of SNP-based demultiplexing may be more broadly applicable for non-human species, especially for different laboratory or wild animals. In particular it is of broad importance to determine if SNP-based demultiplexing works for species in which sample pooling is already a standard practice.

In this project we set out to learn how well SNP-based demultiplexers work in an array of non-human species. To address this we applied souporcell (Heaton et al. 2020), a SNP-based demuxer with minimal requirements for supporting genomic resources, to a variety of species and experimental setups. To benchmark the SNP-based demux results, we compared souporcell results to ground truth cell assignments determined from a range of multiplexing methods. We found that SNP-based demuxers can successfully demultiplex snRNA-seq and scRNA-seq datasets without any prior information about the samples and with only a *de novo* transcriptome as a reference. Based on our analyses, we suggest that SNP-based demuxers can facilitate effective experimental design via the demultiplexing of pooled scRNA-seq data from a vast range of species with few requirements for genomic resources. This further broadens the potential uses of scRNA-seq to study cellular processes in most organisms.

## Results

### Highly accurate SNP-based demultiplexing of *in silico* pooled single cell/nuclei RNA-seq from zebrafish, monkey, and axolotl

We first explored the performance of SNP-based demultiplexing methods when applied to a high quality, low complexity dataset from a well resourced organism, the zebrafish. Two useful tools for applying SNP-based demultiplexing are available for zebrafish, a high quality genome (Howe et al. 2013) a common SNP variant Variant Call Format (VCF) file for the species (LaFave et al. 2014). A recent study collected single animal scRNA-seq datasets from the thymus of zebrafish (Rubin et al. 2022). These data present an opportunity to synthetically pool samples to test SNP-based demultiplexing on data from a non-human species. Though synthetically pooled data are less challenging for a SNP-based demultiplexing algorithm than experimentally pooled samples, the individually sequenced libraries provide a ground truth and enable definitive benchmarking analysis of SNP-based assignments.

After processing each zebrafish scRNA-seq sample individually, we performed *in silico* pooling of three samples (Figure 1A). This synthetic pooling creates a pooled sample that has cell origin information as well as synthetic doublets. The production of synthetic doublets further bolsters the ability to test and validate the accuracy of cell and doublet assignments. The synthetically pooled data was then demultiplexed with the SNP-based demuxer souporcell. Souporcell was chosen as the SNP-based demultiplexer of choice for this study because it has a particularly low minimum threshold for required genomic resources and does not require prior information about individual samples.

**Figure 1:**
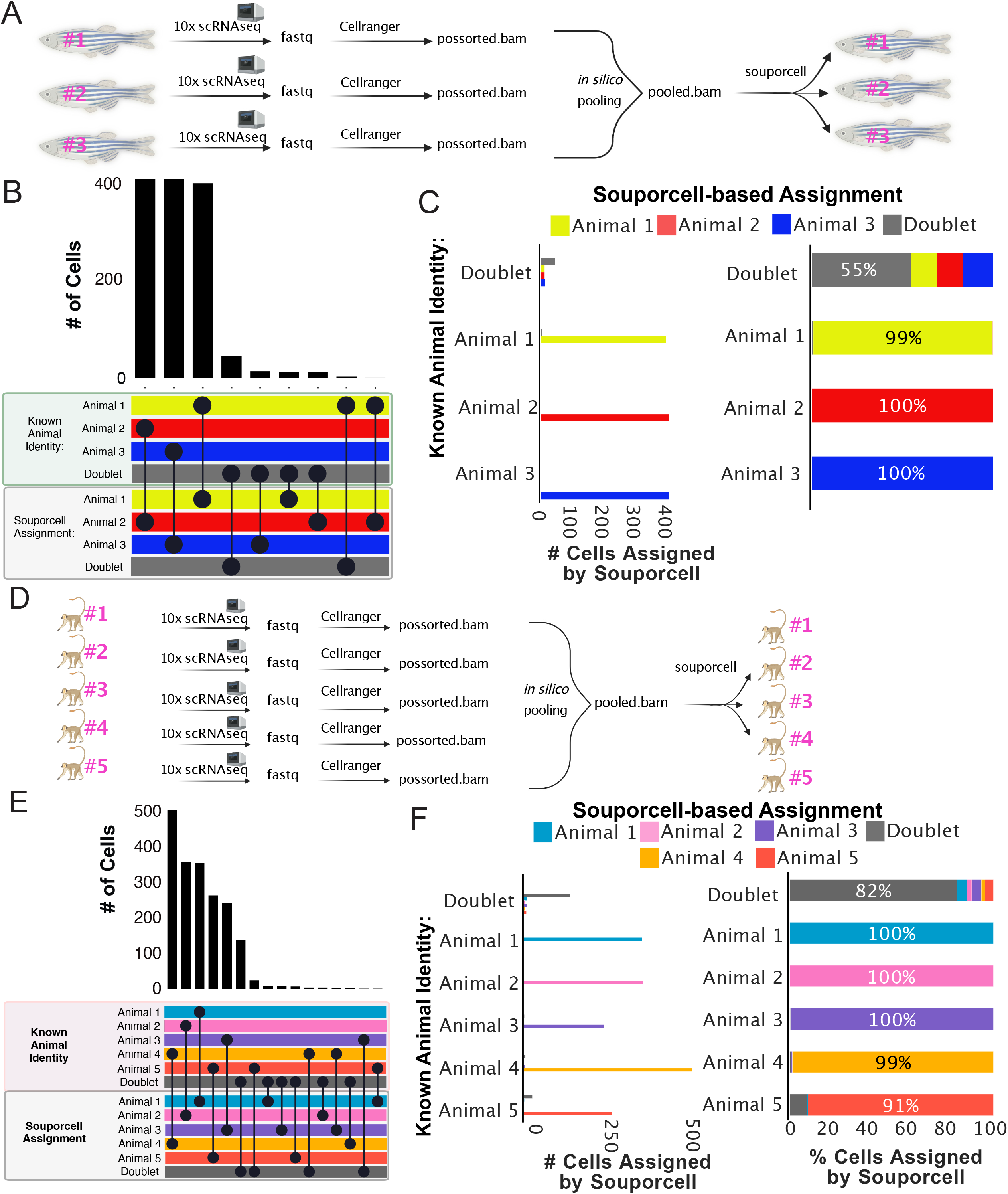
SNP-based demultiplexing enables demultiplexing of synthetically pooled zebrafish and African green monkey scRNA-seq data. A) Conceptual diagram of benchmarking analysis for zebrafish data. B) Upset plot comparing cell assignments by souporcell to known animal identities for each cell. Souporcell assignments were matched with known identities by correlation analysis. C) Bar plots quantifying the distribution of souporcell assignments for cells from each animal. Left: Of cells known to originate from each animal, the number of those cells assigned by souporcell to each animal is plotted. Right: Of cells known to originate from each animal, souporcell assignments are shown as a percentage of total cells assigned to that animal. D-F) Same as A-C but for African green monkey scRNA-seq data.

To analyze the demultiplexing accuracy, SNP-based cell assignments were assessed for correlations with ground truth animal origin. We found a strong agreement between souporcell assignments and ground truth animal identities (Figure 1B-C). We then analyzed how many cells of each known animal source were assigned to each animal by souporcell (Figure 1C, left as total cell quantities, right as percentages). SNP-based demultiplexing of this synthetically pooled zebrafish scRNA-seq data was highly accurate, resulting in correct cell assignments of 99% to 100% of cells based on their known animal origin (Figure 1C). This suggests that genetic demultiplexing is a viable means to enable sample pooling in zebrafish.

In addition to comparing the SNP-based demultiplexing accuracy for identifying the cell source of individual cells, we wondered how well the souporcell doublet detection worked on these samples for heterotypic doublets (i.e., doublets from two genetically distinct individuals). Homotypic doublets created during synthetic pooling were removed, as souporcell relies on intergenotypic doublet detection. We found that souporcell missed almost half of the synthetic heterotypic doublets in the pooled dataset (Figure 1C). The relatively poor performance (55% heterotypic doublets identified) of souporcell at identifying synthetic heterotypic doublets from this zebrafish data made doublet detection the largest discrepancy in this benchmarking analysis. The imperfect doublet detection performance on this dataset could be due to a variety of factors including a lack of read depth for some cells in the synthetic pool. In the Discussion we propose multiple pathways to improve doublet detection in future analyses.

We next evaluated the results of SNP-based demultiplexing when applied to synthetically pooled scRNA-seq data from the African green monkey. The African green monkey is a pre-clinically relevant species, with a published genome (Warren et al. 2015) and SNP variant VCF file readily available for use (Y. S. Huang et al. 2015). Considering previous findings that SNP-based demultiplexers have issues when applied within mouse strains (Mylka et al. 2022), we were curious if SNP-based demultiplexing would work on another non-human mammal. For this experiment, five individually sequenced green monkey scRNA-seq datasets were pooled *in silico* and subsequently demultiplexed (Figure 1D)(Speranza et al. 2021). To facilitate pooling, we selected cells with high gene expression levels for demultiplexing (see Methods). We found that SNP-based demultiplexing correctly identified 91% to 100% of the cells analyzed in the pooled green monkey dataset (Figure 1E-F). Contrasting the zebrafish data, 82% of heterotypic doublets from green monkeys were correctly identified by souporcell. Overall these results suggest that SNP-based demultiplexing can be a highly accurate and efficient method for demultiplexing pooled single cell data from non-human primates.

Finally, we also assessed results of SNP-based demultiplexing of synthetically pooled single nuclei data from a salamander species, the axolotl. The axolotl is an example of the type of organism for which scRNA-seq has enabled cell level study of regeneration and immunology for the first time (Gerber et al. 2018; Lin et al. 2021; Leigh et al. 2018; Rodgers, Smith, and Voss 2020; Lust et al. 2022; Ye et al. 2022). Additionally, a genome(Nowoshilow et al. 2018; Smith et al. 2019) and SNP variant VCF file (Timoshevskaya et al. 2021) are available for the axolotl, but its large genome provides a distinct challenge when using computational tools. snRNA-seq data from the brain of three individual axolotl was recently published (Lust et al. 2022). We used *in silico* pooling for three individually sequenced axolotl snRNA-seq datasets (Supplemental Figure 1A) (Lust et al. 2022). 95% to 99% of the cells from each individual axolotl dataset were correctly demultiplexed from the pool (Supplemental Figure 1B-C). Similarly to the zebrafish dataset, we found that souporcell failed to identify a large proportion of the synthetic heterotypic doublets (Supplemental Figure 1C). These results suggest that SNP-based demultiplexing may accurately demultiplex pooled samples in any single cell compatible organism with genetic heterogeneity.

### SNP-based methods successfully demultiplex experimentally pooled, high sample number, scRNA-seq data from *Xenopus*

Synthetically pooled scRNA-seq data lack challenging aspects of physically pooled scRNA-seq data including the presence of ambient RNA and real heterotypic doublets. Therefore, we were next interested in benchmarking SNP-based demultiplexing accuracy in a more realistic and challenging experimental scenario: experimentally pooled data with a high sample number and without a common SNPs VCF file. We analyzed a published dataset of *Xenopus laevis* scRNA-seq data containing eight experimentally pooled samples from three Xenopus transgenic lines that each overexpress a different fluorescent gene (Lin et al. 2021) (Figure 2A). *Xenopus* are another common laboratory animal that has a published high quality genome (Session et al. 2016). Unlike the synthetically pooled datasets analyzed previously, for this dataset there is no experimental means to determine the empirical cell origin for cells in this dataset (Figure 2A). However, fluorescent gene mRNA detection can be used to assign cells to specific transgenic animal lines, enabling validation of SNP-based demuxing at the level of transgenic line.

**Figure 2:**
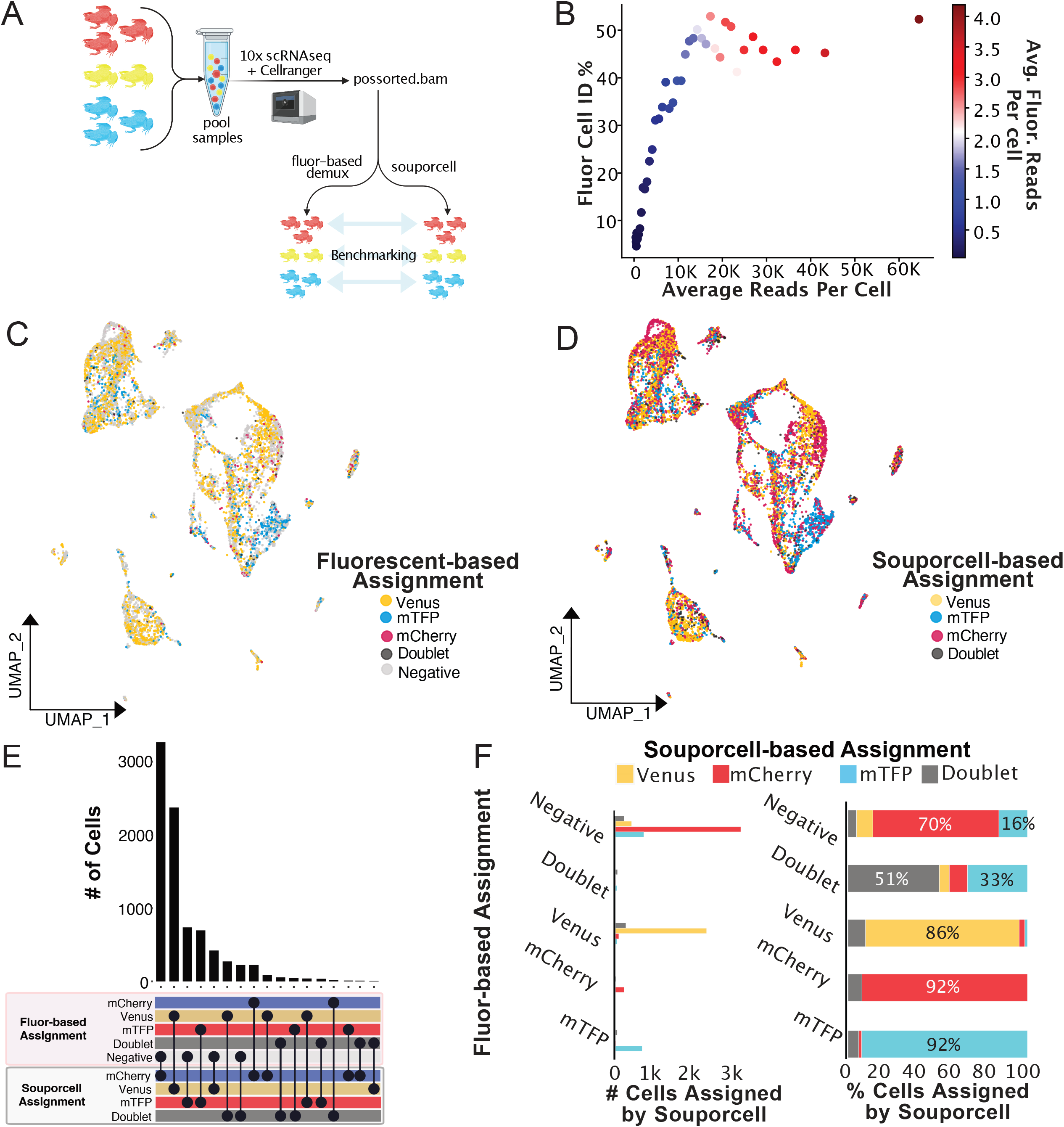
Experimentally pooled, high sample number Xenopus data demultiplexed by SNP-based methods. A) Conceptual diagram of benchmarking analysis for Xenopus data. B) Cell identification percentage (Fluor Cell ID %) by fluorescent based assignment is plotted against average read depth. All cells were sorted by read depth, and binned into 40 groups before calculating Fluor Cell ID%, and average total and fluorescent read depth. Fluor Cell ID % is defined as the percentage of cells in each bin that were assigned to any of the three transgenic animal identities by fluorescent based demultiplexing analysis. Binned data are colored by the average number of summed fluorescent reads per cell. Subsequent analysis plots focused on high accuracy cells with between 5,000 and 40,000 mapped reads, and >0 summed fluorescent gene reads. C) UMAP plot of Xenopus pooled scRNA-seq data colored by fluorescent based cell assignments. D) UMAP plot of Xenopus pooled scRNA-seq data colored by souporcell assignments relabelled according to correlating transgenic animal name. E) Upset plot comparing cell assignments by souporcell to fluorescent based assignments. F) Bar plots quantifying the distribution of souporcell assignments for cells from each animal. Left: Of cells assigned to each transgenic line through fluorescent-based assignments, the number of those cells assigned by souporcell to each category is plotted. Right: Of cells assigned to each transgenic line through fluorescent-based assignments, souporcell assignments are shown as a percentage of total cells in that category.

To identify the transgenic line origin based on fluorescent mRNA counts, we co-opted the MULTIseqdemux(McGinnis et al. 2019) algorithm to assign donor identities based on the transgenic line expressed fluorescent gene counts. Though this approach succeeded in assigning the *Xenopus* transgenic line of origin, the low number of fluorescent gene counts left many cells without sufficient data to make an assignment prediction (Supplemental Figure 2, Figure 2B). This low detection of fluorescent gene transcripts is a common problem when using fluorescent marker genes for demuxing pooled data (Lin et al. 2021) and underscores the importance of alternative means to demultiplex pooled data. To avoid benchmarking results with low quality cells, a stringent filtering cutoff was used to ensure that only cells likely to have accurate origin assignment by the fluorescent gene-based demultiplexing method would be used to benchmark SNP-based demultiplexing.

We first assessed the filtered *Xenopus* data for correlation between fluorescent and SNP-based cell assignments. We observed a remarkable similarity in demultiplexing the eight animal dataset with these two orthologous methods (Figure 2C-E, Supplemental Figure 2). To further quantify the souporcell assignment accuracy, we evaluated how souporcell performed in comparison to transgenic line assignments made by fluorescent based-assignment (Figure 2F). We found that the two methods agree well, as demonstrated by the souporcell assignment agreement on 86% to 92% of cells in each sample (Figure 2F). Further, a large number of cells were assigned as “Negative” by the fluorescent-based approach, which could then be computationally “rescued” and assigned a transgenic line by souporcell. This is particularly notable for the mCherry expressing cells, which were difficult to assign by the fluorescent-based assignment method (Figure 2C-D). Also of interest, we found that one of the eight transgenic animals was almost completely lacking from the dataset, only displaying dozens of total cells. Though individual animal assignments by souporcell could not be validated, this suggests animal dropout for the dataset that would not be identified without demultiplexing.

Approximately 6% to 9% of cells from each transgenic line assigned by fluorescent-based assignments were called doublets by souporcell. It is not clear from the available data if these are true doublets being identified only by souporcell, or if the SNP-based demuxer is over-assigning doublets. However, assuming the worst case scenario: that souporcell is over-assigning doublets, this is a relatively harmless mistake considering that a standard analysis would subsequently remove these doublets. Overall these results display a high degree of accuracy for SNP-based assignments in a complex, experimentally pooled mixture of *Xenopus* cells.

### Successful SNP-based demuxing of pooled fluorescent Pleurodeles samples without a genome or previous SNP information

After observing that SNP-based demultiplexing was reliable on various commonly used model species with genomes and with or without common SNP VCF files, we wanted to test the limits of what computational resources are required for accurate SNP-based demultiplexing. We chose to use only a *de novo* transcriptome without common SNP VCF files. The Spanish ribbed newt, *Pleurodeles waltl*, is an emerging regenerative model organism, which has a high quality *de novo* transcriptome (Matsunami et al. 2019) but no available common SNPs VCF file. We first set out to assess souporcell demux assignments on pooled splenocytes from three transgenic *Pleurodeles* newts, which express different fluorescent proteins under the same ubiquitous promoter (CAG)(Joven et al. 2018; Eroglu et al. 2022). We designed this experiment to only contain non-erythroid spleen cells from one individual of each transgenic newt line (Supplemental Figure 5), making it technically feasible to benchmark souporcell cell assignments for individuals through comparisons to fluorescent-based assignments (Figure 3A).

**Figure 3:**
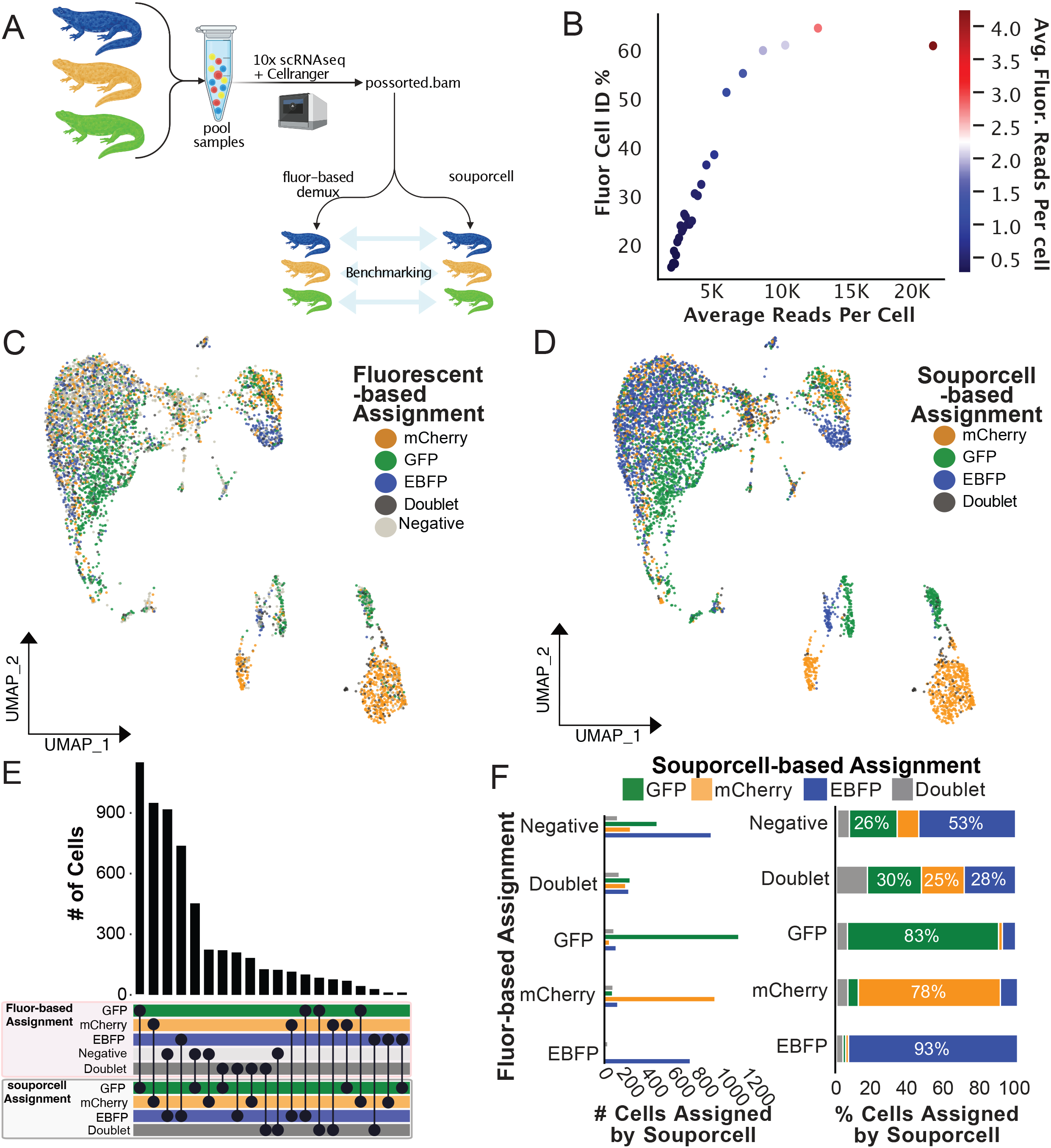
Experimentally pooled Pleurodeles scRNA-seq from fluorescently expressing transgenic animals accurately demultiplexed by souporcell. A) Conceptual diagram of benchmarking analysis for this Pleurodeles dataset. B) Cell identification percentage (Fluor Cell ID %) by fluorescent based assignment is plotted against average read depth. All cells were sorted by read depth, and binned into 40 groups before calculating Fluor Cell ID%, and average total and fluorescent read depth. Fluor Cell ID % is defined as the percentage of cells in each bin that were assigned to any of the three transgenic animals by fluorescent based demultiplexing analysis. Binned data are also colored by the average number of summed fluorescent reads per cell. Subsequent analysis plots focused on high accuracy cells with between 5,000 and 40,000 mapped reads, and >0 summed fluorescent gene reads. C) UMAP plot of Pleurodeles pooled scRNA-seq data colored by fluorescent-based cell assignments. D) UMAP plot of Pleurodeles pooled scRNA-seq data colored by souporcell assignments relabelled according to correlating transgenic animal line. E) Upset plot comparing cell assignments by souporcell to fluorescent based assignments. F) Bar plots quantifying the distribution of souporcell assignments for cells from each animal. Left: Of cells assigned to each transgenic animal through fluorescent-based assignments, the number of those cells assigned by souporcell to each animal is plotted. Right: Of cells assigned to each transgenic animal through fluorescent-based assignments, souporcell assignments are shown as a percentage of total cells in that category.

As done in the *Xenopus* analysis,we selected only cells that had sufficient read depth and fluorescent gene detection for benchmarking (Figure 3B). We found that fluorescent-based and souporcell assignments show a high degree of correlation (Figure 3C-D, Supplemental Figure 3). The fluorescent-based approach assigns many cells as “Negative” (63% of cells pre-filter, 29% of cells after filtering) (Figure 3B-D). Further, even though these samples were pooled and sequenced as one sample, we find dramatic variation between individuals in the *Pleurodeles* splenocyte data (i.e., clusters derived from only one animal). The heterogeneity of sample representation in different cell clusters highlights the need for demultiplexing of pooled scRNA-seq data. Without demultiplexing, erroneous conclusions on novel cell states or types may arise.

We found a high degree of agreement between fluorescent-based cell assignments and SNP-based assignments from *Pleurodeles* scRNA-seq data (Figure 3E). Of cells assigned by fluorescent-based demuxing to one of the three transgenic animals, 78% to 93% of those cells were correctly identified by souporcell (Figure 3F). When the two methods disagree, the most prevalent occurrence is “Negative” fluorescent-based cell assignments that souporcell assigned to one of the transgenic animals (i.e., the “rescue” we also found in the *Xenopus* data). Similar to the *Xenopus* dataset, we attribute these fluorescence-based “Negative” assignments to the low capture of fluorescent gene reads (Figure 3B-C, Supplemental Figure 3). The next most common discrepancy between the two demultiplexing approaches were cells labeled as “Doublet” by fluorescent-based demuxing, for which souporcell disagrees. The low number of mapped fluorescent reads in these samples means that a “Doublet” assignment by fluorescent-based demuxing likely indicates that a particular cell had one to two counts of two distinct fluorescent genes. This could be indicative of a doublet, but it is not certain that these cells are missed doublet assignments because fluorescent-based doublet assignments too often rely on little biological data.

### SNP-based demultiplexing succeeds in two-species, pooled single cell RNA-seq as shown by benchmarking against lipid hashing

We next wanted to determine whether SNP-based demultiplexers could succeed in demuxing scRNA-seq datasets containing cells pooled from multiple species. The ability to demultiplex pooled single cell sequencing experiments from multiple species would be particularly useful for cross-species analyses and ecological studies. This approach would capitalize on the so-called “barnyard” approach that is typically used when investigating doublet rates in new single cell technologies. For this experiment we pooled splenocytes from two *Pleurodeles waltl*, and two *Notophthalmus viridescens* (Supplemental Figure 4A). *Notophthalmus* is another salamander species studied for its regenerative capacity but equipped with only a *de novo* transcriptome and no published common SNP VCF file. To investigate SNP-based assignments and doublets in this dataset, we applied a barnyard style analysis to this dataset and found a general agreement between these two approaches (Supplemental Figure 4A-G).

Within this experiment of dual species pooled 10X scRNA-seq libraries, distinct lipid hashing cellplex multiplexing oligo (CMO) labels were applied to label the origin of each of the *Pleurodeles* samples, and one hashing label was used for the pooled *Notophthalmus* samples, totalling three hashing labels (Figure 4A). This approach provided a means to benchmark souporcell against the current best practice and the only available commercial multiplexing strategy for cells from species without specific hashing antibodies. For the analysis, sample identity determined by applying MULTIseqDemux to CMO data was used to evaluate SNP-based demultiplexing results. We constructed and aligned to a dual species index made from the superTranscriptome (Davidson, Hawkins, and Oshlack 2017; Abdullayev et al. 2013; Matsunami et al. 2019) for each of these species. We removed low quality, low depth cells for further benchmarking analysis and identified which souporcell individual assignments corresponded to each CMO through analyzing the correlation of assignments between the two methods (Figure 4B). One important note, it appeared that many cells with high CMO reads per cell have relatively low biological reads per cell (Figure 4B), suggesting that while these cells can be demultiplexed they would not provide relevant biological information.

**Figure 4:**
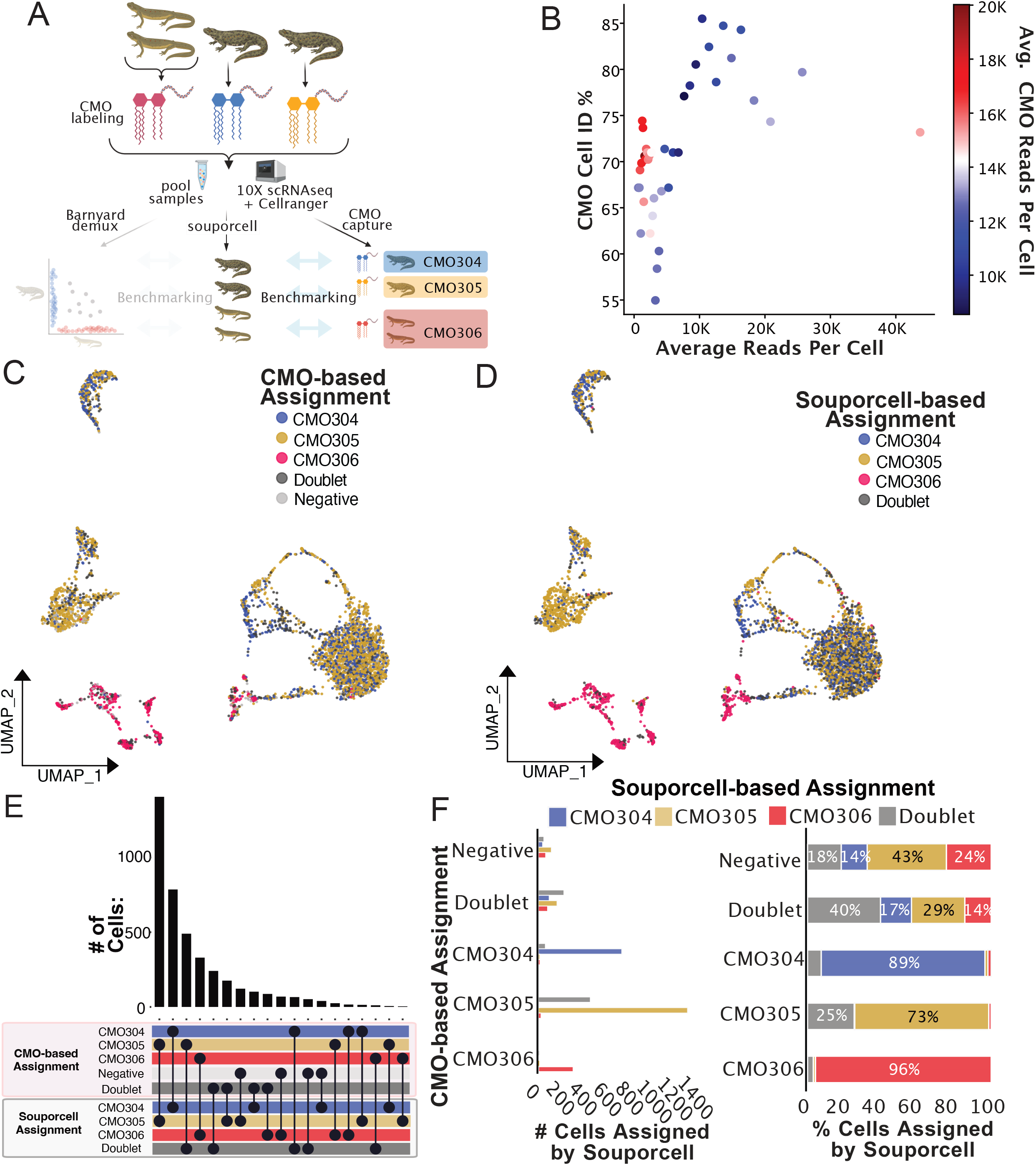
Lipid linked CMO-based demultiplexing of two salamander species, four animal pooled scRNA-seq dataset further confirm accuracy of souporcell results. A) Conceptual diagram of benchmarking analysis for this multi-species dataset. B) Cell identification percentage (CMO Cell ID %) by CMO-based assignments is plotted against average read depth. All cells were sorted by read depth, and binned into 40 groups before calculating CMO Cell ID%, and average total read depth per cell. CMO Cell ID % is defined as the percentage of cells in each bin that were any of the three CMO groups via CMO analysis. Subsequent analysis plots focused on high accuracy cells with between 5,000 and 40,000 mapped reads. C) UMAP plot of Pleurodeles and Notophthalmus pooled scRNA-seq data colored by CMO assignments. D) UMAP plot of Pleurodeles and Notophthalmus pooled scRNA-seq data colored by souporcell assignments relabelled according to correlating CMO labels. E) Upset plot comparing cell assignments by souporcell to CMO-based assignments. F) Bar plots quantifying the distribution of souporcell assignments for cells from each CMO group. Left: Of cells assigned to each CMO group, the number of those cells assigned by souporcell to each identity is plotted. Right: Of cells assigned to each CMO group, souporcell assignments are shown as a percentage of total cells in that category.

We found that souporcell sample assignments correlated tightly with CMO-based assignments (Figure 4C-E). When CMO-based assignments were interpreted as ground truth, the accuracy of souporcell assignments was clear, with 73% to 96% agreement between the two methods for assignment to the three CMO-based assignments (Figure 4F). The largest difference in assignments between these methods was that souporcell identified many doublets in the data from one CMO labeled sample. One possible explanation for this disagreement is imperfect CMO application leading to spillover labeling of other samples. Overall, these analyses demonstrate high agreement between CMO-based and SNP-based demultiplexing of this data from a notably complex experiment with species with poor genomic resources.

## Discussion

The commercial appearance of single cell sequencing technologies has enabled the study of complex tissues from any species. The technical and financial hurdles posed by these technologies can discourage their use, especially when biological replicates are needed to produce reliable datasets. We demonstrate that sample pooling is a useful and viable tool for empowering reproducible single cell datasets. It is critical that pooling and demultiplexing approaches are also species agnostic, as one of the major advantages of scRNA-seq is its broad applicability across organisms.

The results presented in this study are in strong support of using sample pooling and SNP-based demultiplexers in scRNA-seq studies in any species which possesses between individual genetic variability. We show here in five species with varying genomic resources that the demuxer souporcell produced highly accurate demuxing results. We found that the ground truth afforded by *in silico* pooled samples was particularly important for thorough primary validation of SNP-demultiplexing accuracy in non-human species. However, for benchmarking the accuracy in more realistic but challenging, complex experimental setups, the lack of similarly perfect ground truth method led us to utilize a diverse swath of methods, each with unique strengths. Importantly, no experimentally-driven demultiplexing approach is expected to be mistake free, which means that some loss of accuracy attributed here to souporcell is likely due to the approaches being used for validation. For species in which it is unclear whether SNP-based demultiplexers will work successfully, we recommend a pilot experiment in which a secondary demultiplexing method, or *in silico* pooling, is used to benchmark SNP-based demux results.

Animal to animal variability is expected in many tissues, especially in the immune system for animals that encounter different pathogens. This poses problems for studies without replicates or those that do not demultiplex pooled datasets. Demultiplexing of pooled scRNA-seq is required to associate metadata with individual samples for batch correction, differential expression analysis, and confirmation of sample representation across scRNA-seq datasets and cell clusters (Squair et al. 2021). The potential for heterogeneity in sample representation across cell clusters can be clearly seen in Figure 3, where some cell clusters are clearly driven by one or two of the three individuals present in the pool. Unlike some model species like mice, which are often kept in clean conditions to make immune perturbations uniform, the aquatic environment in which anamniote species are housed may produce heterogeneity in the immune experiences for each individual. However, the use of biologically variable samples in single cell sequencing studies requires the sequencing of higher numbers of biological replicates in order to capture sufficient representation of natural variability. These studies include animals like ‘dirty’ or outbred mice (Beura et al. 2016), lab-reared aquatic organisms, or wild animals (i.e., ecoimmunology), thus requiring pooling techniques to control study costs and batch effects. Though a variety of batch correction algorithms are available for single cell studies (Tran et al. 2020), these cannot be used if the metadata distinguishing distinct replicates is not associated with pooled data. Based on our findings in this study, we suggest the use a SNP-based demultiplexing approach to separate pooled single cell samples, even if only to confirm that all individuals added to a pool are represented in the final results.

An additional benefit of pooled single cell experiments is that it enables the superloading of cells followed by heterotypic doublet detection. Our synthetic pooled data results saw varied success in heterotypic doublet detection, and therefore do not currently argue strongly in favor of superloading samples. In fact, the most consistent problem with souporcell assignments noted in this study is that the souporcell internal doublet detection may not be sensitive enough in some cases. Mistakes in doublet detection were particularly noticeable in the *in silico* pooled demultiplexing experiments with zebrafish and axolotl data when using default doublet detection settings. The production and use of an improved SNP VCF file for any given species could improve heterotypic doublet detection by souporcell. The internal doublet detection method in souporcell, troublet, can also be adjusted to optimize doublet detection. If run separately from the souporcell pipeline, troublet has customizable thresholds for doublet and singlet detection that may enable more accurate heterotypic doublet calling. Additionally, multiple independent programs designed specifically for doublet detection using transcriptomic instead of genotypic information are available for scRNA-seq data including DoubletDetection (Gayoso and Shor 2022), Solo (Bernstein et al. 2020), and Scds (Bais and Kostka 2020). We expect that the optimized application of internal or external doublet detection algorithms along with improved bioinformatic resources for each species, would improve doublet detection and facilitate superloading pooled single cell data for use with SNP-based demultiplexing in non-human species.

Many bioinformatic tools for scRNA-seq are not easily applied to all species. In this project we circumvent the need for genomic resources like a high quality, well annotated genome, or a population wide common SNP genotypes VCF file. These resources are not available for most species. In the case of souporcell, we found that these two resources are not required since this program can be applied to datasets with only a *de novo* transcriptome and no VCF input. A secondary hurdle for applying SNP-based demuxers is that these tools often struggle when applied to datasets from species with large reference genomes or *de novo* transcriptomes (which typically have many contigs). When using *de novo* transcriptomes (i.e., *Pleurodeles* or *Notophthalmus*) we modified the default souporcell pipeline to enable remapping of reads (Supplemental Table 1, see Methods). Though these steps should not be necessary for most species, they were helpful in enabling the use of souporcell with a *de novo* transcriptome reference and may be useful for the many species which do not have reference genomes available.

Overall we successfully applied SNP-based methods to demultiplex pooled single cell data from multiple species including *in silico* pooled scRNA-seq and snRNA-seq data from zebrafish (Rubin et al. 2022) African green monkey (Speranza et al. 2021), and axolotl (Lust et al. 2022) respectively, as well as experimentally pooled scRNA-seq data from *Xenopus* (Lin et al. 2021), *Pleurodeles*, and experimentally pooled dual salamander species scRNA-seq. Our benchmarking results suggest that SNP-based demultiplexing in these species is accurate relative to other available demultiplexing approaches. We hope that this study will increase awareness of single cell pooling and SNP-based demultiplexing approaches for research communities not yet using these methods. Including SNP-based demuxers in experimental designs for future (and past) studies will greatly expand single cell-based discoveries. This will facilitate work in well-known and lesser studied species by lowering the financial and technical hurdles of producing adequately powered single cell experiments. We predict that both species agnostic, and cross-species comparative studies are going to be increasingly fruitful in uncovering biological insights and the application of SNP-based demultiplexing with minimal genomic resources is critical for future research.

## Materials and Methods

### Animal handling and ethics

All experiments were carried out in post-metamorphic *Pleurodeles waltl* and *Notophthalmus viridescens* at Karolinska Institutet and were performed according to local and European ethical permits. *Notophthalmus viridescens* and *Pleurodeles waltl* were raised in-house. All animals were maintained under standard conditions of 12-hour light/12-hour darkness at 18-24°C (Joven, Kirkham, and Simon 2015). Prior to all experiments, animals were fully narcotized in 0.1% tricaine in housing water. For *Pleurodeles* in the fluorescent pool experimental animals were housed in carbon supplemented filtered tap water (55g Tetra Marine SeaSalt, 2 teaspoons Ektozon N Salt, 2.5mL of water conditioner/dechlorinator (Seachem Prime - Vattenberedningsmedel), 20mL of Yokuchi Bitamin - Multivitamin and 10mL of calcium supplementation (Easy-Life Calcium) into 100L of water). For *Pleurodeles* and *Notophthalmus* in the Cellplex experiment animals were housed in the water as described but modified to only have sea salt, Ektozon, and calcium solution.

### Collection of splenocytes for scRNAseq

#### Fluorescent pooling experiment

Spleens were harvested from three separate *Pleurodeles waltl* and processed as individual samples in parallel. All animals were post-metamophic newts from established transgenic lines close to sexual maturity: one female tgTol2(CAG:Nucbow CAG:Cytbow)^Simon^ (5.67g weight, 10.8cm snout-to-tail length) (Joven et al. 2018), one male tgSceI(CAG:loxP-GFP-loxP-Cherry)^Simon^ (5.36g, 11.1cm)(Joven et al. 2018), and female tgTol2(CAG:loxP-Cherry-loxP-H2B:YFP)^Simon^ (6.25g, 10.8cm)(Eroglu et al. 2022). Forceps and iridectomy scissors were used to remove the spleen in one piece, making sure that the forceps did not tear the spleen. Iridectomy scissors were used to carefully remove connective tissue. A 70μm nylon mesh filter was inserted into a 50mL conical and 1mL of ice cold 0.7X PBS was added to pre-wet the filter. The spleen was placed on the pre-wetted filter and slowly mashed through the filter using the plunger stopper end of a 3mL syringe. Once the spleen appeared translucent, the plunger and filter were thoroughly washed with ~10mL ice cold 0.7X PBS and making sure no PBS was left on the plunger or filter. The 0.7X PBS solution was then poured into a 15mL conical and centrifuged at 300g for 5 min at 4°C in a swinging bucket rotor. Supernatant was decanted and cells were resuspended in 1mL of 0.7X PBS. Cells were then counted using trypan blue to assess viability. Viabilities of non-erythroid cells (based on cellular morphology) were 85%, 98.4% and 91.6% in eBFP, eGFP, and mCherry animals, respectively. Fluorescent activated cell sorted was used to sort for fluorescent positive cells (Figure 5A-D). The samples have been analyzed without any removal of red blood cells using an INFLUX (BD) cytometer; the selected cells were sorted using a 100 μm nozzle in bulk into 1.5 ml Eppendorf tubes. The cell preparations and the sorted cells were kept in 4 °C throughout the sorting. The results were analyzed with FlowJo software 10.8.1.

Debris and erythrocytes (note: erythrocytes do not express fluorescent markers under the CAG promoter and are far larger than other splenocytes (Supplemental Figure 5E) were excluded with the gating strategy in the side scatter-forward scatter and singlet discrimination plots revealing separated fluorescent subpopulations of GFP, mCherry and BFP as confirmed by sorting on microscopy slides (Supplemental Figure 6). We sorted the subpopulations with the highest expression levels of each fluorescent tag. In total, 4×10^5^ cells of GFP+, 4×10^5^ of mCherry+ and 3.15×10^5^ BFP+ were isolated in individual 1.5mL eppendorf tubes. GFP and mCherry expression was marked, but BFP expression was dim. 500μL of each solution was then added to an individual 1.5mL eppendorf for the fluorescent pool sample (trial mixture of pooled cells shown Figure 5D). This was centrifuged in an eppendorf at 4°C at 300g on a tabletop centrifuge. The supernatant was carefully removed and resuspended in the remaining volume. Cells were then manually counted and adjusted to a concentration targeting the collection of 10,000 cells on the 10x Genomics controller.

#### Cellplex/barnyard pooling experiment

Spleens from one female *Notophthalmus viridescens* (4.45g and 10.6cm snout-to-tail) and one male *Notophthalmus viridescens* (3.55g and 10.2cm) were collected, pooled into one tube, and then processed as described in “*Fluorescent pooling experiment*” excluding FACS cell sorting. The only modifications to the above described processing is that cells were kept at room temperature throughout and that cells were resuspended in 0.7X PBS in 0.04% ultrapure BSA.

For *Pleurodeles waltl*, spleens were removed from animals as described in “*Fluorescent pooling experiment*” from one adult tgSceI(CAG:loxP-GFP-loxP-Cherry)^Simon^ female (23.5g and 16.1cm snout-to-tail length), one male tgSceI(CAG:loxP-GFP-loxP-Cherry)^Simon^ (13.95g and 15.7cm) animal. *Pleurodeles* were processed as individual samples. After the spleen was thoroughly mashed through the pre-wetted 70μm nylon filter and the filter was washed with 10mL of 0.7X PBS the cells were centrifuged at 300g for 5 min. Splenocytes were resuspended cells in 1mL of sterile filtered 1X ACK (http://cshprotocols.cshlp.org/content/2014/11/pdb.rec083295.short) to lyse red blood cells. After one minute of lysis, cells were diluted with 10mL of 0.7X PBS and filtered through a 70μm nylon mesh filter and centrifuged at 300g for 5 min. Cells were then resuspended 0.7X PBS in 0.04% ultrapure BSA.

The pooled *Notophthalmus* sample and the individual *Pleurodeles* samples were then taken through the 10x Genomics 3’ Cellplex labeling protocol (Demonstrated Protocol, CG000391) the only modifications being the use of 0.7X PBS + 0.04% BSA for all wash and resuspension steps. Samples were stained with CM304 (*Pleurodeles* female), CMO305 (*Pleurodeles* male), and CMO306 (pool of *Notophthalmus* samples, one male and one female). Samples were manually counted and pooled at equal ratios immediately prior to loading onto the 10× Genomics Chromium Controller targeting 9,000 cells in total.

### Preparation and sequencing of single-cell RNA sequencing libraries

Chromium single cell 3’ kit v3 (10× Genomics) was used according to the manufacturer’s instructions.

### Generation of SuperTranscriptomes and corresponding gtf files

A *Pleurodeles waltl* de novo transcriptome from (Matsunami et al. 2019) was downloaded from (https://figshare.com/articles/dataset/Trinity_Pwal_v2_fasta_gz/7106033/1), and unzipped. The Trinity (Haas et al. 2013) singularity image v2.11.0 was then used to generate a *Pleurodeles waltl* Super Transcriptome like so:

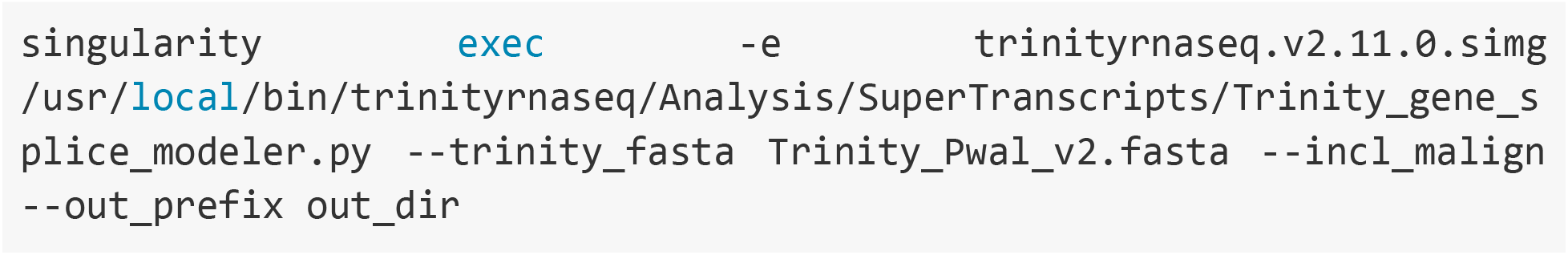

A *Notophthalmus* de novo transcriptome from (Abdullayev et al. 2013) was downloaded from (https://sandberglab.se/static/data/papers/redspottednewt/reference_transcripts_v2.fa.gz) and decompressed. Transcripts were clustered with cd-hit-est (W. Li and Godzik 2006; Fu et al. 2012)

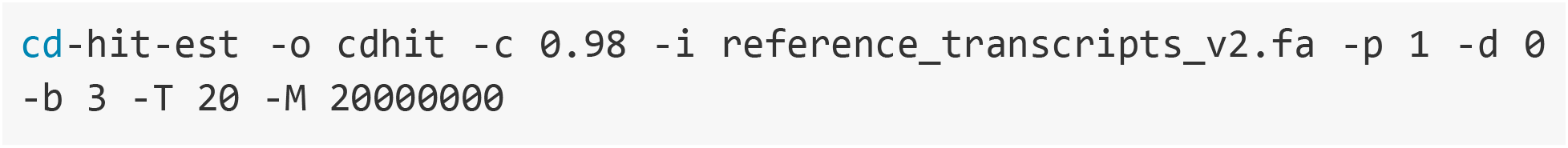

The resulting cdhit.clstr file was then parsed using clstr2txt.pl from the cdhit package to generate a clusters.txt file. The id and clstr columns were obtained from this file and used to make a info.clusters.txt file to use with Lace(Davidson, Hawkins, and Oshlack 2017). Lace v1.14.1 was run: Lace_run.py reference_transcripts_v2.fa info.clusters.txt -t --cores 16 -o Noto_superTrans. The resulting gff files from both superTranscriptomes were converted to a gtf using AGAT perl script agat_convert_sp_gff2gtf.pl (Dainat et al. 2022).

### Zebrafish souporcell demultiplexing

Fastq files from a previously published paper (Rubin et al. 2022) (SRA accessions: SRR17218111, SRR17218113, SRR17218114, SRR17218091, SRR17218092) were downloaded from SRA using prefetch followed by fasterq-dump with flags split-files and include-technical. Files were renamed to 10X fastq format (e.g., sample_S1_L001_I1_001.fastq.gz) and then aligned using Cell Ranger v7.0.0 count to GRCz11 with the corresponding gtf for GRCz11 filtered via Cell Ranger mkgtf for protein_coding genes.

To merge bams in silico a vcf file was downloaded from https://research.nhgri.nih.gov/manuscripts/Burgess/zebrafish/downloads/NHGRI-1/danR_er11/danRer11Tracks/NHGRI1.danRer11.variant.vcf.gz (LaFave et al. 2014) and subsequently filtered using bcftools filter --include ‘MAF>=0.05’ and then sorted using bcftools sort -Oz. The chromosomes between the vcf and gtf did not match so bcftools annotate --rename-chrs was used to change chromosome names in the vcf (e.g., chr1 to 1).

The sample specific bam outputs were then merged using Vireo’s synty_pool.py script as following:

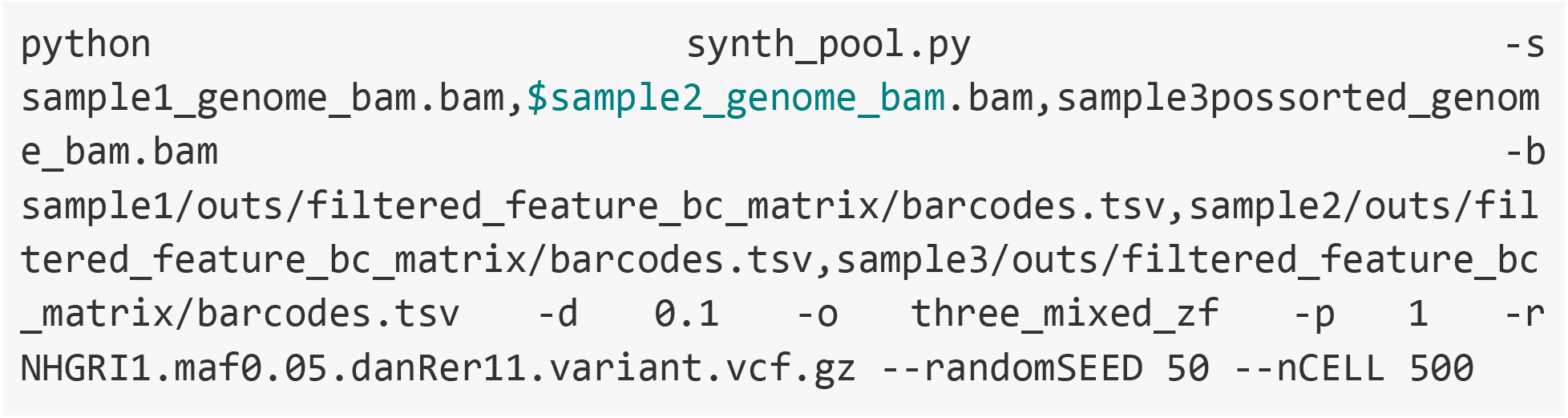

Souporcell was run using souporcell_pipeline.py with inputs: merged BAM output from synth_pool.py, the output barcodes_pool.tsv from synth_pool.py, the genome fasta (Danio_rerio.GRCz11.dna.primary_assembly.fa), N = 3, and vcf file NHGRI1.maf0.05.danRer11.variant.vcf.gz

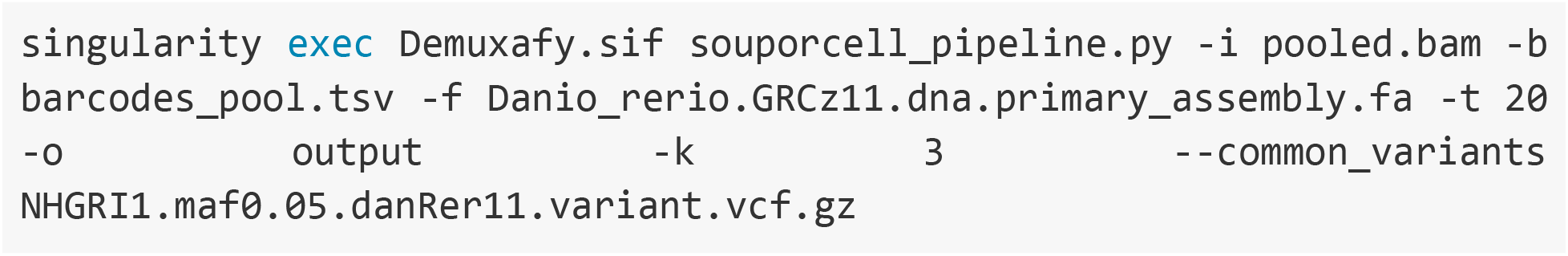

### Xenopus souporcell demultiplexing

The Cellranger BAM file (https://sra-pub-src-2.s3.amazonaws.com/SRR13600554/107606_Xen_Pool_BL7_10_14dpa.bam.1) from a previously published publicly available dataset(Lin et al. 2021) of fluorescent expressing *Xenopus laevis* cells pooled from 8 animals: two blastemas of two siblings (CAGGs:Venus), three blastemas of three siblings (CAGGs:mCherry, B51), 3 samples from 3 siblings (CAGGs:TFPnls, G48) was download directly. samtools(H. Li et al. 2009) was use to index the BAM prior to running default souporcell pipeline with the barcodes file (https://ftp.ncbi.nlm.nih.gov/geo/samples/GSM5057nnn/GSM5057661/suppl/GSM5057661_107606_Xen_Pool_BL7_10_14dpa_barcodes.tsv.gz), *Xenopus laevis* genome fasta (https://sra-pub-src-2.s3.amazonaws.com/SRR13600553/Xenbase_v9.2.fa.1), and N = 8. Note: this specific reference includes the plasmid sequences necessary for mapping to fluorescent sequences.

### Axolotl in silico mixing and souporcell demultiplexing

Fastq files from a previously published (Lust et al. 2022) axolotl single nucleus RNA sequencing data set were downloaded. Libraries from ArrayExpress (E-MTAB-11638) labeled “reseq” and from samples D_1, L_1, and M_1, three individual animals all run on individual wells on a 10x chip were downloaded. To make a Cellranger reference the axolotl genome (AmexG_v6.0-DD) was downloaded from https://www.axolotl-omics.org/dl/AmexG_v6.0-DD.fa.gz along with a gtf (AmexT_v47-AmexG_v6.0-DD.gtf) https://www.axolotl-omics.org/dl/AmexT_v47-AmexG_v6.0-DD.gtf.gz, which required the removal of white space (i.e, sed ‘s/\ \[/_/g’) for use with Cellranger (v7.0.0) mkref. Cellranger count was run on each library individually resulting in three position sorted BAM files from samples D_1, L_1, and M_1. BAM files were merged using synth_pool.py from Vireo(Y. Huang, McCarthy, and Stegle 2019) using the vcf downloaded from (http://ambystoma.uky.edu/hubExamples/hubAssembly/hub_AmexG_v6/AmexG_v6.hub_data/SNP_vcf_tracks/ddMale_to_AmexGv6.vcf.gz) which was filtered using BCFtools v1.11

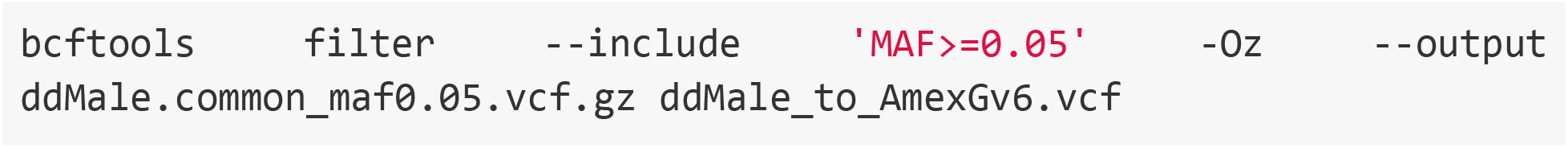

and then sorted:

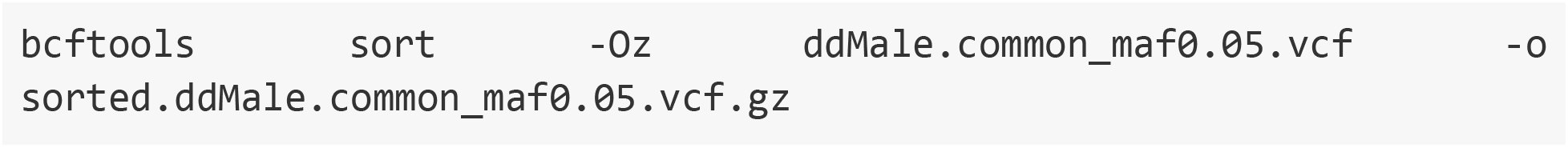

Barcodes.tsv files were obtained from filtered outputs of Cellranger count for each library. Doublet rate (-d) was set to 0.1 and --randomSEED 50

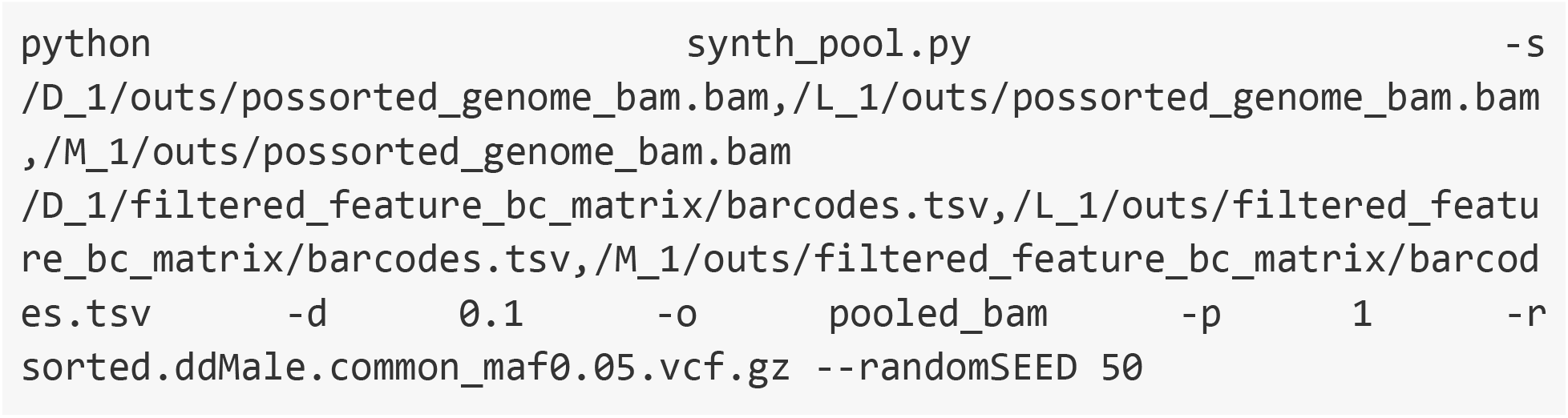

Note: we only expected the troublet portion of souporcell to be capable of detecting heterotypic doublets, so for downstream analysis of this synthetically pooled data, we removed all homotypic doublets.

The pooled BAM was indexed using samtools index -c which made a .csi index and was renamed to .bai for use in souporcell. Souporcell was run using souporcell_pipeline.py with inputs: merged BAM output from synth_pool.py, the output barcodes_pool.tsv from synth_pool.py, the genome fasta (AmexG_v6.0-DD.fa), N = 3, vcf file ddMale.common_maf0.05.vcf.gz, and --skip_remap SKIP_REMAP.

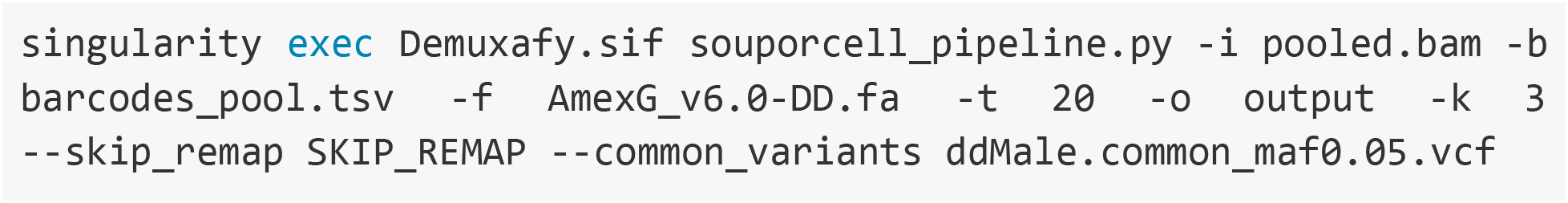

### Green Monkey (Chlorocebus aethiops) in silico mixing and souporcell demultiplexing

Five samples were downloaded from a previously published dataset(Speranza et al. 2021). Data were downloaded from SRA using prefetch followed by fasterq-dump with flags split-files and include-technica. Fastq files were obtained from AGM1_Medialstinal Lymph Node (SRR12507774-SRR12507781), AGM3_Medialstinal Lymph Node (SRR12507790-SRR12507797), AGM5_Medialstinal Lymph Node (SRR12507806-SRR12507813), AGM7_Medialstinal Lymph Node (SRR12507822-SRR12507829), and AGM9_Medialstinal Lymph Node (SRR12507846-SRR12507853). *Chlorocebus aethiops* has a robust VCF file available (European Variation Archive: PRJEB7923) that needs to be used in conjunction with genome assembly Chlorocebus_sabeus 1.1 (GCA_000409795.2). This assembly did not have an annotation file available and we generated a gtf file for this GenBank assembly using minimap2 (H. Li 2018) and the below described steps.

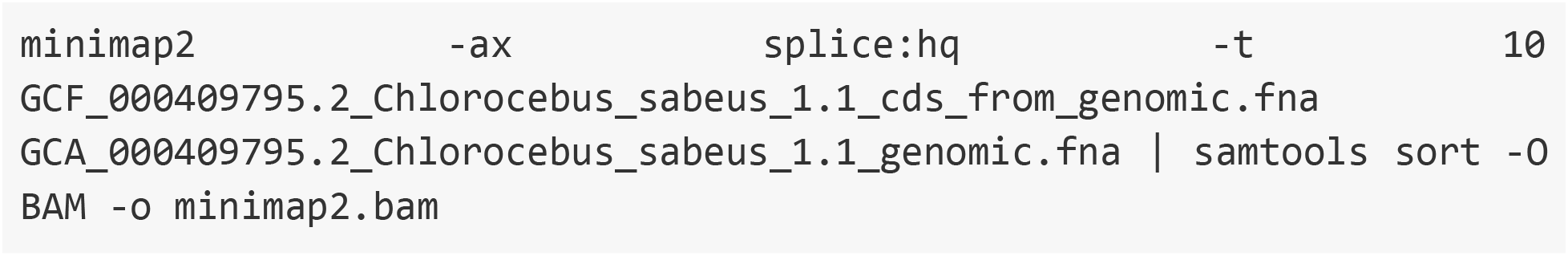

Followed by bedtools (Quinlan and Hall 2010):

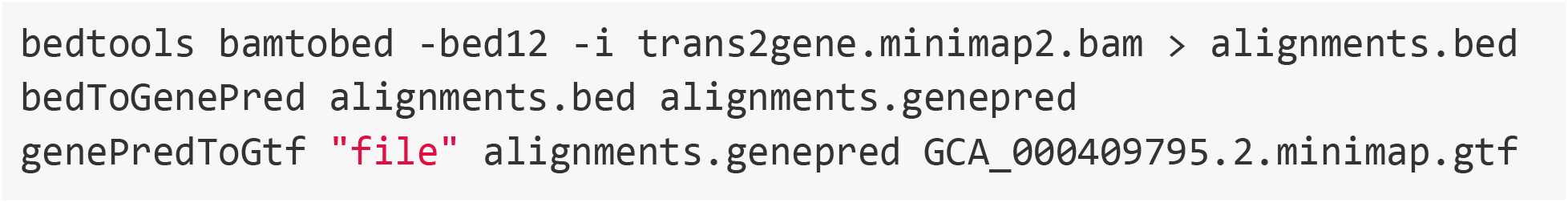

This gtf was then used with GCA_000409795.2_Chlorocebus_sabeus_1.1_genomic.fna to generate a cellranger reference using cellranger mkref. Cellranger count was used with default settings to align the downloaded libraries. To merge the BAM outputs we firsted used bcftools(Danecek et al. 2021) to filter, sort, and change chromosomes name in the vcf:

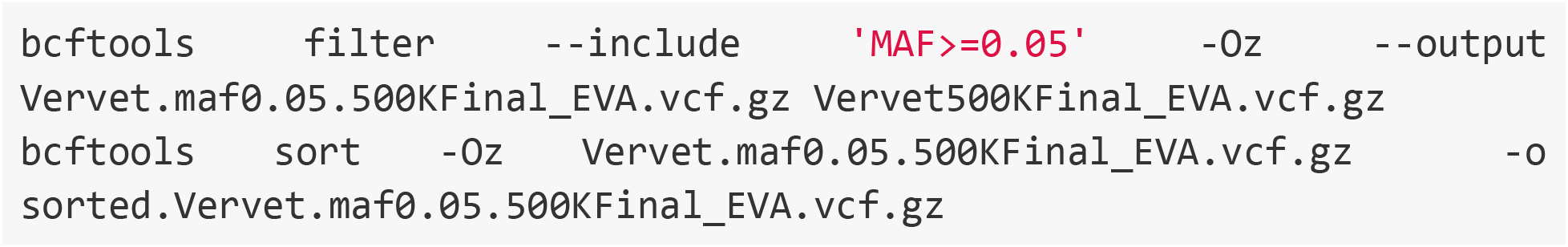

And then converted chromosome names in this vcf to match the chromosomes in the bam files after mapping. chr_name_conv.txt has the format of “1 CM001941.2”, with 1 being the original chromosome number and CM001941.2 being the chromosome accession number; this was done from chromosome 1 to 29.

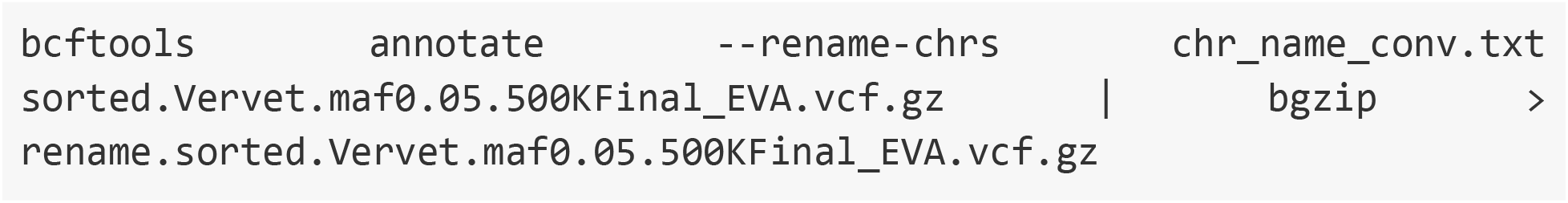

Due to low reads/cell in these libraries, we selected barcodes using Seurat with between 1000 and 2000 features. These filtered barcode files were used with 10× subset-bam (https://github.com/10XGenomics/subset-bam) (e.g., subset-bam --bam possorted_genome_bam.bam --cell-barcodes filtered.barcodes.tsv --out-bam filtered.bam) to create BAMs with these high quality cells. BAMs were subsequently merged using Vireo:

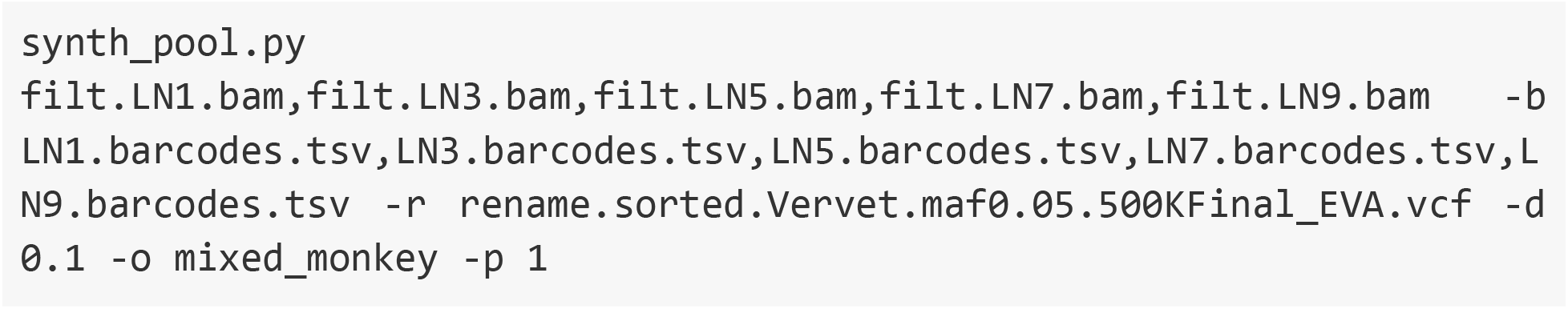

Finally, cells were SNP-demultiplexed using souporcell:

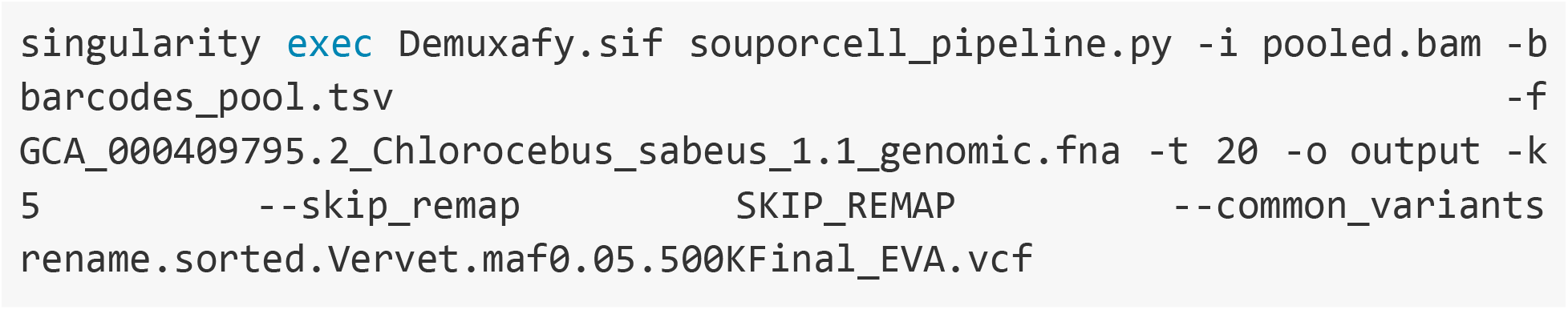

### Pleurodeles mapping and SNP-based demultiplexing

Cellranger 7.0.0 mkref command was used with the above listed Pleurodeles superTranscriptome fasta and gtf files to produce a Cellranger compatible reference. Cellranger 7.0.0 count command was then used to map and count reads over the transcriptome for the three transgenic animal scRNA-seq dataset.

Souporcell (Heaton et al. 2020) related processes were all run from a demuxafy (Neavin et al. 2022) singularity image (image version 1.0.3). The remapping and variant calling stages of souporcell were run externally due to problems with timeouts on the remapping process with the large salamander transcriptome, and issues with the souporcell internal freebayes command failing. The VCF from freebayes was then used in souporcell pipeline with the --skip_remap SKIP_REMAP and--common_variants ${VCF}. Full scripts used for souporcell processes are included below. Summary of souporcell run details can be found in Supplemental Table 1.

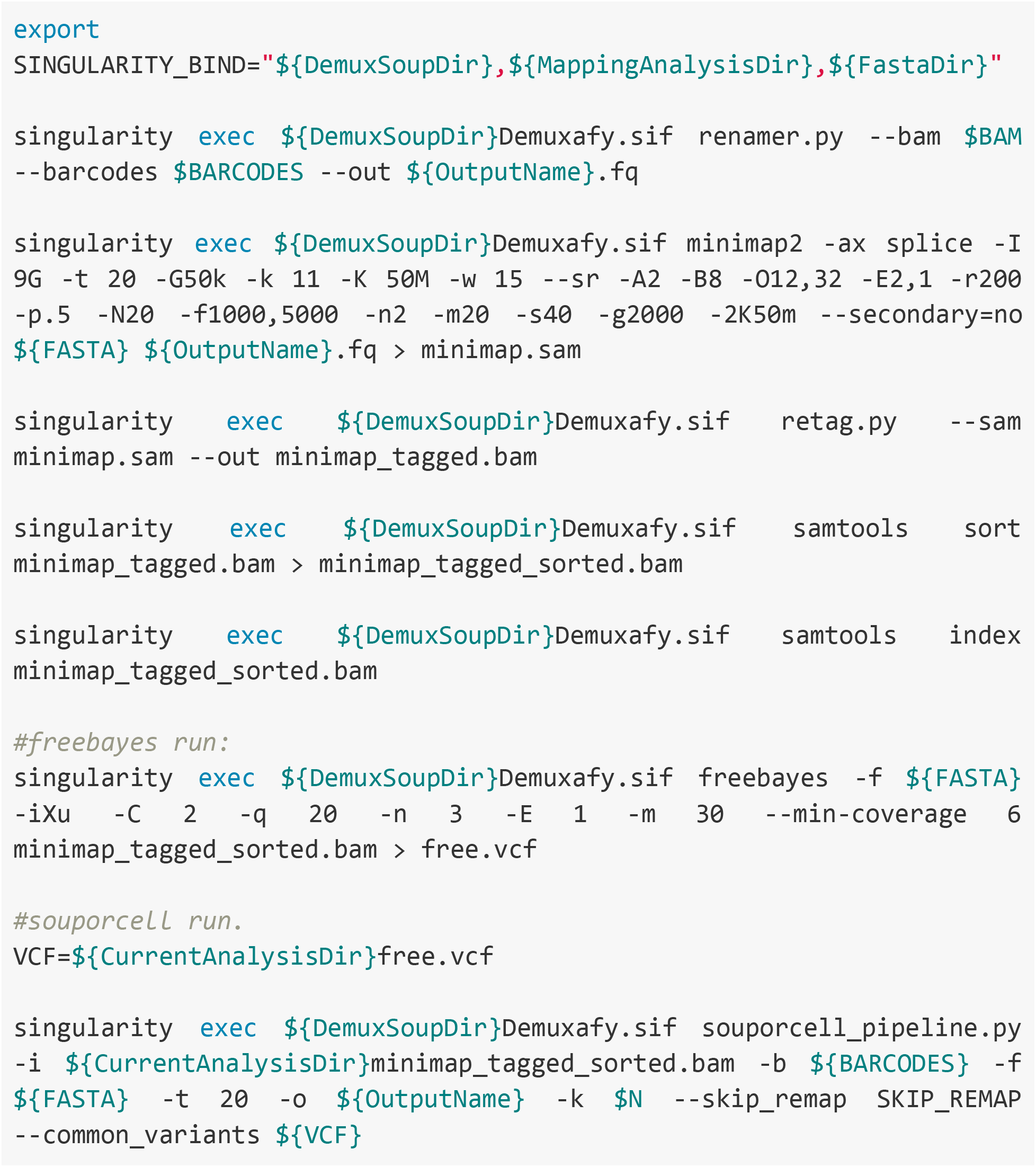

### Pooled Pleurodeles and Notophthalmus mapping and SNP-based demultiplexing

A dual species Cellranger 7.0.0 reference was made utilizing the superTranscriptomes and corresponding gtf files (described above) from the two species using Cellranger mkref command. The two species, four animal, pooled scRNA-seq from Pleurodeles and Notophthalmus dataset was then mapped to this dual species index using

Cellranger 7.0.0 count command. Additionally, the Cellranger 7.0.0 Multi command was used to assess multiplexing Cellplex information for all cells in the same dataset. For the Cellranger Multi command, the following flags were used: (expect cells 10000, min-assignment-confidence 0.6). Souporcell demultiplexing was run identically to above on the Pleurodeles only samples but with N=4, and the relevant fastq and dual species reference transcriptome fasta files.

### Souporcell analysis details summary

A summary table reviews the computational details used to run Souporcell on the above datasets (Supplemental Table 1). Overall the standard souporcell default pipeline was used initially, but additional flags or pieces of the souporcell pipeline were run externally when this failed due to memory constraints or other problems. Souporcell was run from a demuxafy (Neavin et al. 2022) singularity container (image version 1.0.3). SNP numbers in each VCF were counted using : grep “##” VCFname | wc -l

For analyses of *Pleurodeles* and dual species datasets, the first two steps of the souporcell pipeline were run separately and then the output from these was introduced back into the souporcell pipeline for completion (Supplemental Table 1). This allowed us to adjust computational parameters in the remapping stage that permitted the function to finish.

### Analysis and benchmarking of souporcell assignments. R analysis

A two part analysis in R and then python was used to evaluate the efficacy of souporcell demultiplexing for each dataset. Scripts in R (version 4.1.2) primarily using Seurat (version 4.1.0) (Hao et al. 2021) were used to evaluate souporcell cell assignments found in the clusters.tsv file through comparison with known cell identities or cell assignments based on fluorescence, barnyard analysis, or CMO labeling. Full R scripts for all analyses are deposited in the github page for clarity (https://github.com/RegenImm-Lab/SNPdemuxPaper). Seurat was used to import and analyze single cell gene expression data for all datasets and to analyze the multiplexing capture data for the dual species Cellplex (CMO) labeled dataset. **Cell Filtering:** For UMAP plots and all bar plots in benchmarking analysis cells for all experimentally pooled datasets were filtered prior to analysis to select for cells most likely to have accurate calls by the respective benchmarking assay. Cellranger default filtered cells were parsed to select cells with between 5,000 and 40,000 mapped reads for each dataset. Additionally, for Figures 2 and 3 we removed cells lacking counts of any of the three fluorescent mRNAs. Cell assignments based on fluorescence or Cellplex labeling were made through the analysis of fluorescent reads or Cellplex CMO reads by the Seurat MULTISeqDemux function (autoThresh=T) (McGinnis et al. 2019). Though the MULTIseqDemux function is written to assign cell identities based on CMO labels, we found that it works well with data from overexpressed fluorescent gene mRNAs. For analysis purposes, souporcell demultiplexing numerical cell labels were adjusted to relevant sample names based on correlation analyses to each relevant benchmarking demultiplexing method. UMAP (McInnes, Healy, and Melville 2018) and upset plots (Lex et al. 2014) were generated in R scripts and annotated in Affinity Designer. A dataframe with compiled cell assignment information was exported to python for further analysis due to ease of use.

Session info including package numbers for R analyses are embedded in the github page (https://github.com/RegenImm-Lab/SNPdemuxPaper), and included R version 4.1.2 (2021-11-01).

### Analysis and benchmarking of souporcell assignments. Python analysis

Analyses of read depth versus Cell ID% and quantitative benchmarking of souporcell demultiplexing results was carried out in google colab notebooks shared bellow, using python version 3.7, Numpy version 1.21.6 (Harris et al. 2020), Pandas version 1.3.5 (Reback et al. 2021), Matpotlib version 3.2.2 (Hunter 2007), Scipy version 1.7.3 (Virtanen et al. 2020), Seaborn version 0.11.2 (Waskom 2021).

Google colab links for analyses are located here: Xenopus: https://colab.research.google.com/drive/1lO4ny8Uv9n1lPIbHmZKFPyxWI7gjmZPs?usp=sharing

Pleurodeles:

https://colab.research.google.com/drive/1Zbbpi1WwfKGwrrFhuecrHE3lSjP0Pzsz?usp=sharing

Pleurodeles and Notophthalmus cellplex:

https://colab.research.google.com/drive/12ZNvvfiUt3DL6UpTg8BjQMSd4Y-6yM3u?usp=sharing

Pleurodeles and Notophthalmus barnyard:

https://colab.research.google.com/drive/1JS8kRUGAioDM2IvYa4oBDzNMKLxBhHV_?usp=sharing

Zebrafish, axolotl, and green monkey:

https://colab.research.google.com/drive/1yXzE3WJ05hEJKdy7owiXOpCjYvUm4DDJ?usp=sharing

### Calculation of Cell ID %

All cells were sorted by total mapped read depth, and binned into 40 groups before calculating Cell ID%, and average total and fluorescent read depth where relevant. Cell ID % is defined as the percentage of cells in each bin that were assigned an individual sample identity (non doublet, non negative result) by a demultiplexing method. Cell ID value for each bin was then plotted against average total mapped reads. For datasets including fluorescent transgenic lines, binned data are also colored by the average number of summed mapped fluorescent reads per cell.

Bar plots: Filtered datasets were subset by the animal or animal group assignment from each demultiplex method being used to benchmark souporcell results. Within those subsets, the total cell quantity of cells assigned to each identity by souporcell was plotted (left plots). Alternatively, within each benchmarking demux result subset, the percentage of cells assigned to each identity by souporcell were calculated by dividing by total cells assigned to that identity by the benchmarking demuxer, and multiplied by 100.

## Author Contributions

JFC conceptualized the project, wrote the first draft, and performed computational analysis. AJ helped to generate the scRNA-seq libraries and edited the manuscript. AS provided resources and edited the manuscript. NDL conceptualized the project, generated salamander scRNA-seq data, edited the manuscript, performed computational analysis, supervised the project, and provided resources.

## Acknowledgements

We thank the Eukaryotic Single Cell Genomics core at SciLifeLab for the generation of 10X libraries of Pleurodeles and Notophthalmus samples. Sequencing was performed at NGI Genome Center which is funded by RFI/VR and Science for Life Laboratory, Sweden. The computations, storage and data handling were enabled by resources provided by LUNARC which is part of the Swedish National Infrastructure for Computing (SNIC) and is partially funded by the Swedish Research Council through grant agreement no. 2018-05973. We thank Tomás Pires de Carvalho Gomes and Ashley Maynard for graciously sharing their data with us early and Tomás Pires de Carvalho Gomes for providing feedback on this manuscript. We also thank Garrett Dunlap for critical review and feedback. In addition, we thank the Simon lab for insightful discussions and help with animal care. Diagrams with animals were generated with Biorender.com.

## Funding sources

NDL receives funding from the Knut and Alice Wallenberg Foundation and the Swedish Research Council (Registration # 2020-01486). AS receives funding from European Research Council, Swedish Research Council, and Cancerfonden.

## Data Availability

Data are available on ArrayExpress with accession E-MTAB-12186 for three animal pooled Pleurodeles splenocyte scRNA-seq and ArrayExpress accession E-MTAB-12182 for four animal pooled *Pleurodeles* and *Notophthalmus* splenocyte scRNA-seq. All code used to analyze the data are present in the Methods, in linked Colab notebooks, or via GitHub (https://github.com/RegenImm-Lab/SNPdemuxPaper) All other data used in the manuscript were from previously published works of which accessions are noted in the methods.

## Notes

### Competing Interest Statement

The authors have declared no competing interest.

