## Supplemental_information for "Accurate genotype-based demultiplexing of single cell RNA sequencing samples from non-human animals"

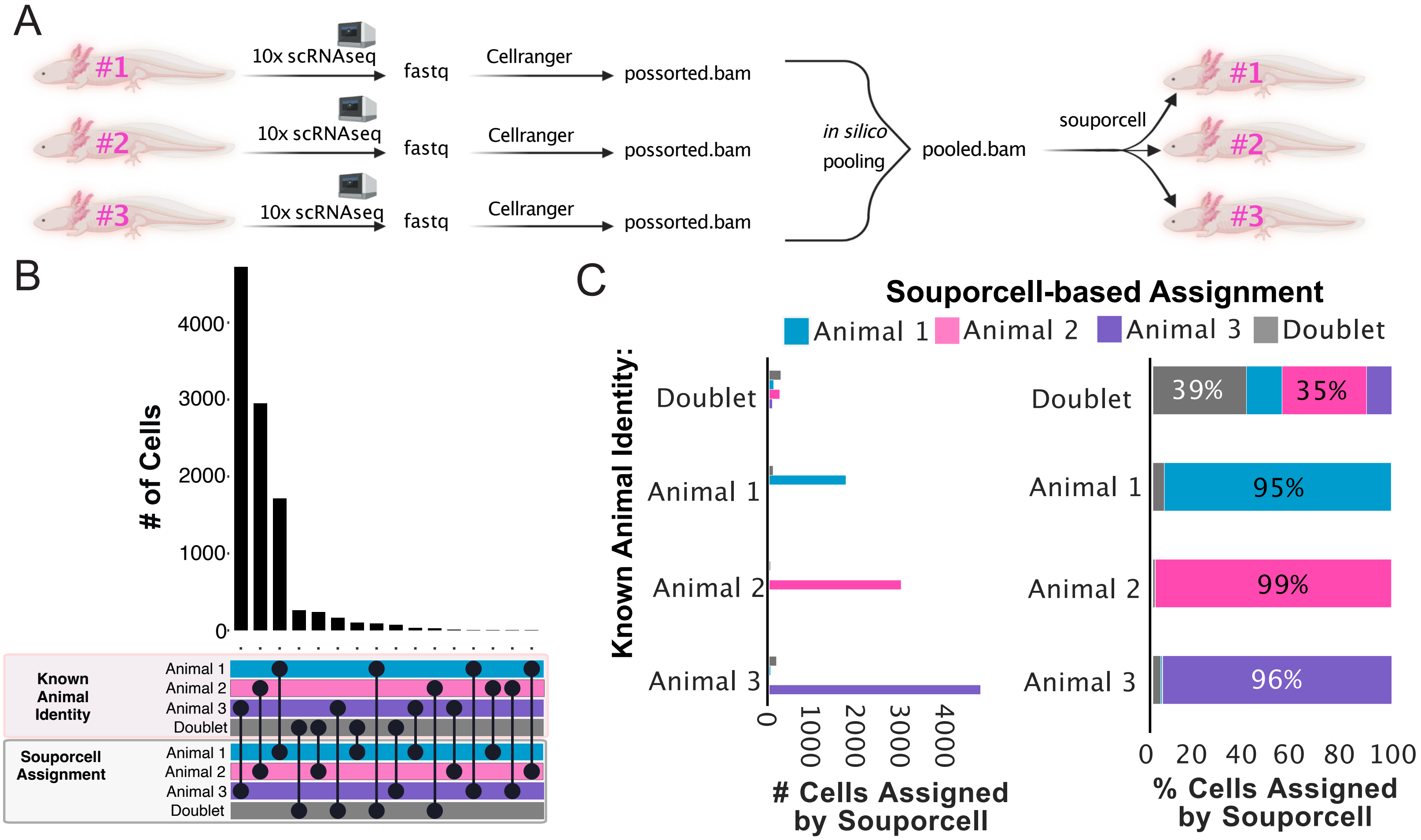


**Supplemental Figure 1**: SNP-based demultiplexing enables demultiplexing of synthetically pooled axolotl snRNA-seq data. A) Conceptual diagram of benchmarking analysis for axolotl data. B) Upset plot comparing cell assignments by souporcell to known animal identities for each cell. Souporcell assignments were matched with known identities by correlation analysis. C) Bar plots quantifying the distribution of souporcell assignments for cells from each animal. Left: Of cells known to originate from each animal, the number of those cells assigned by souporcell to each animal is plotted. Right: Of cells known to originate from each animal, souporcell assignments are shown as a percentage of total cells assigned to that animal.

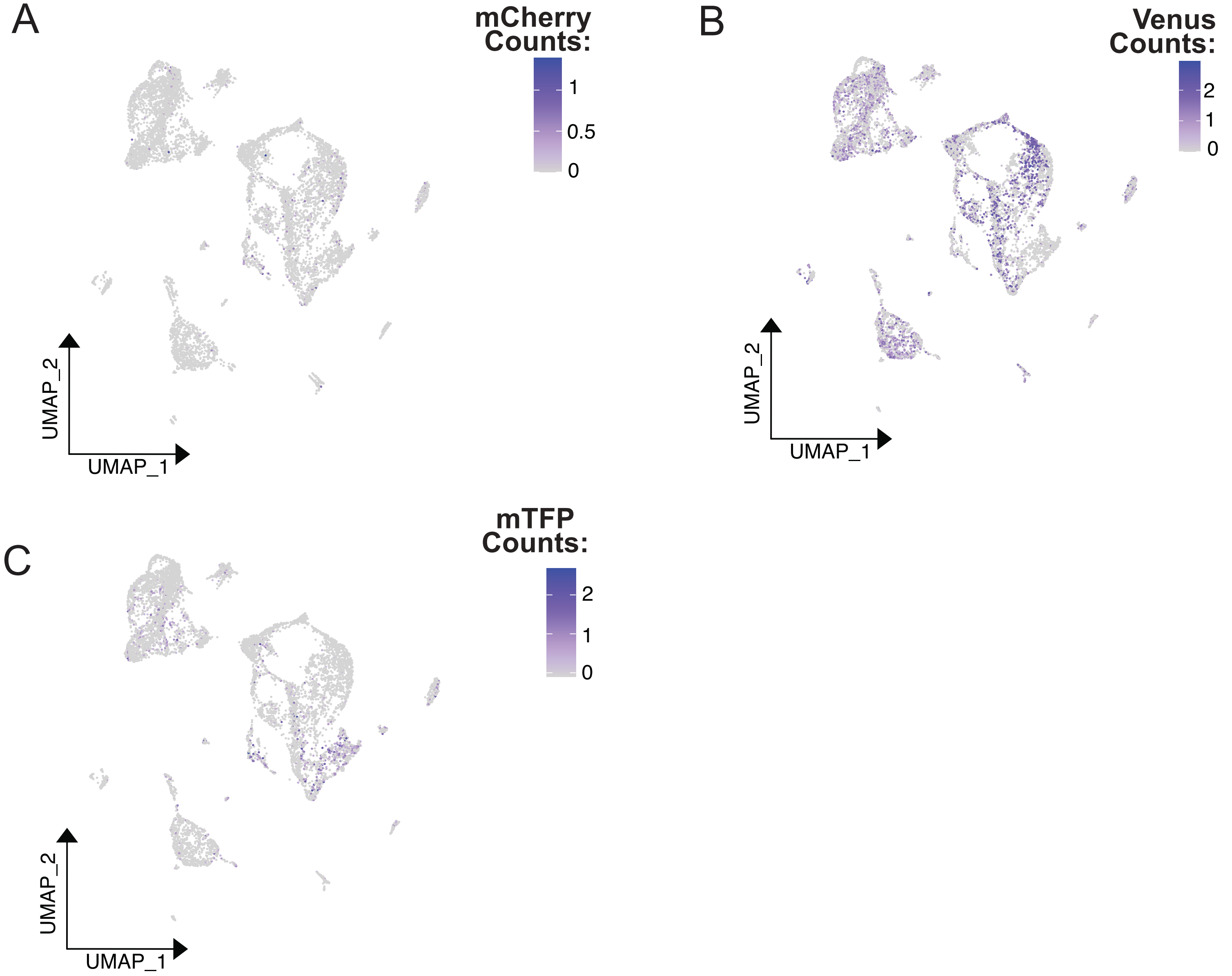


**Supplemental Figure 2:** UMAP plot of pooled Xenopus scRNA-seq data colored by normalized, scaled counts of fluorescent transcripts detected per cell. A) UMAP colored by mCherry Counts. B) UMAP colored by Venus counts. C) UMAP colored by mTFP counts.

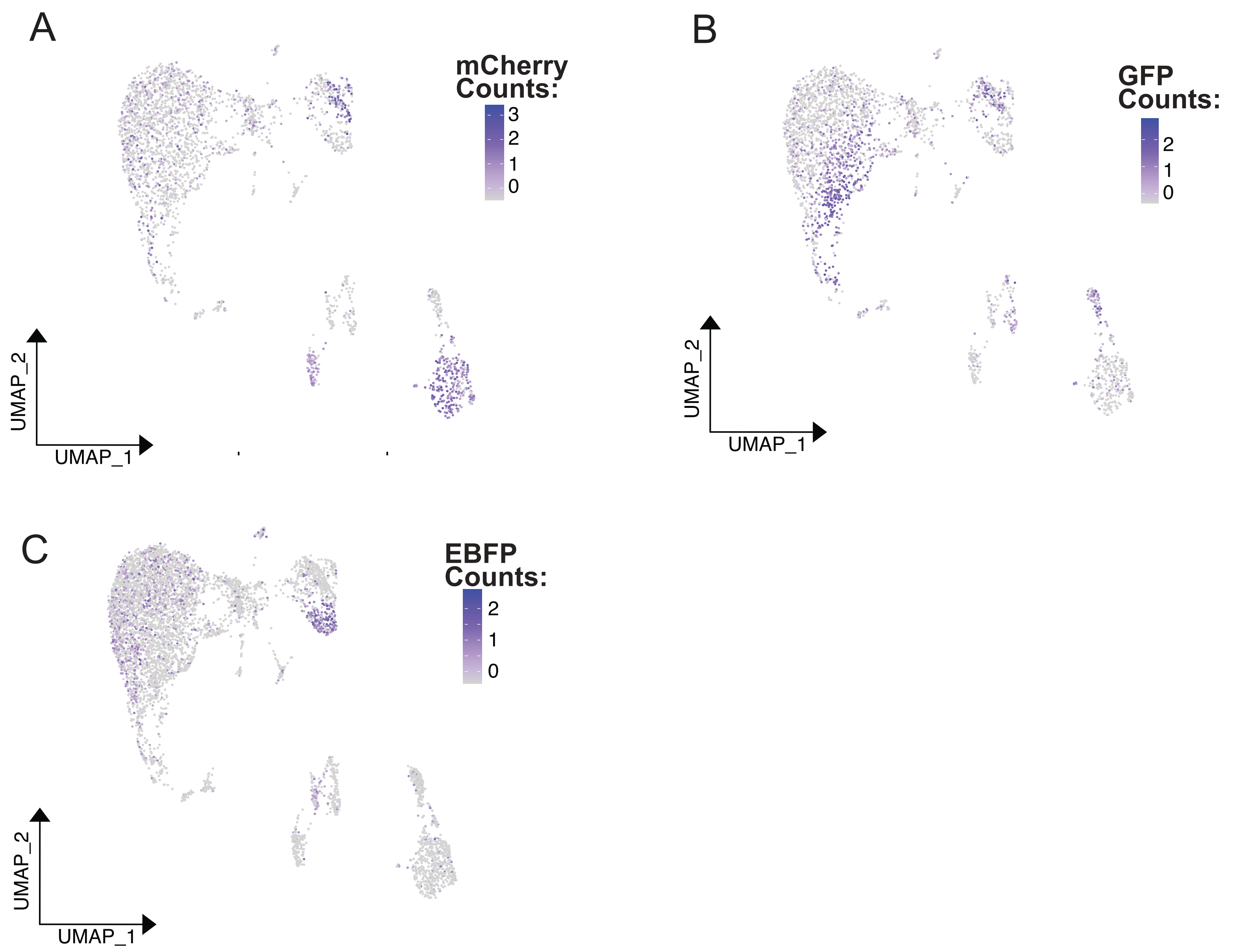


**Supplemental Figure 3:**UMAP plot of Pleurodeles pooled scRNA-seq data colored by normalized, scaled counts of fluorescent transcripts detected per cell.

A) UMAP colored by mCherry Counts. B) UMAP colored by GFP counts. C) UMAP colored by EBFP counts.

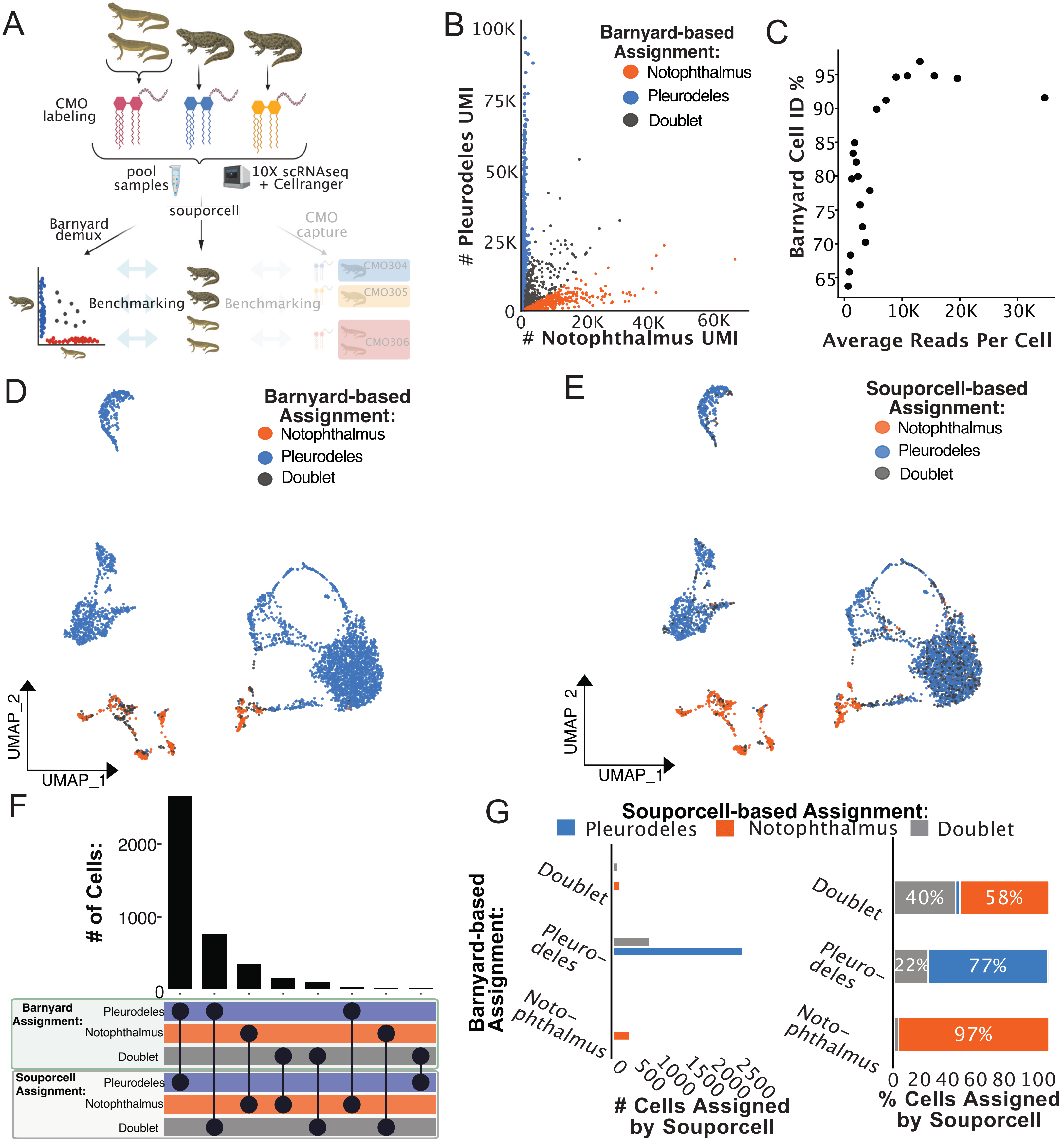


**Supplemental Figure 4:** High agreement between barnyard and souporcell demultiplexing of two species, four animal pooled salamander scRNA-seq dataset. A) Conceptual diagram of experiment and benchmarking analysis for this dataset. B) Barnyard analysis plot revealing the quantity of reads mapping to the transcriptome of each salamander species, and the subsequent barnyard-based cell assignments. C) Cell identification percentage (Barnyard Cell ID %) by barnyard analysis based assignment is plotted against average read depth. All cells were sorted by read depth, and binned into 40 groups before calculating Barnyard Cell ID%, and average total read depth per cell. Barnyard Cell ID % is defined as the percentage of cells in each bin that were assigned to either species identity via barnyard analysis. Subsequent analysis plots focused on high accuracy cells with between 5,000 and 40,000 mapped reads. D) UMAP plot of Pleurodeles and Notophthalmus pooled scRNA-seq data colored by species assigned by barnyard analysis. E) UMAP plot of Pleurodeles and Notophthalmus pooled scRNA-seq data colored by souporcell assignments relabelled according to correlating barnyard determined species. F) Upset plot comparing cell assignments by souporcell to barnyard-based assignments. G) Bar plots quantifying the distribution of souporcell assignments for cells from each animal. Left: Of cells assigned to each species by barnyard-based assignments, the number of those cells assigned by souporcell to each animal is plotted. Right: Of cells assigned to each species through barnyard assignments, souporcell assignments are shown as a percentage of total cells in that category.

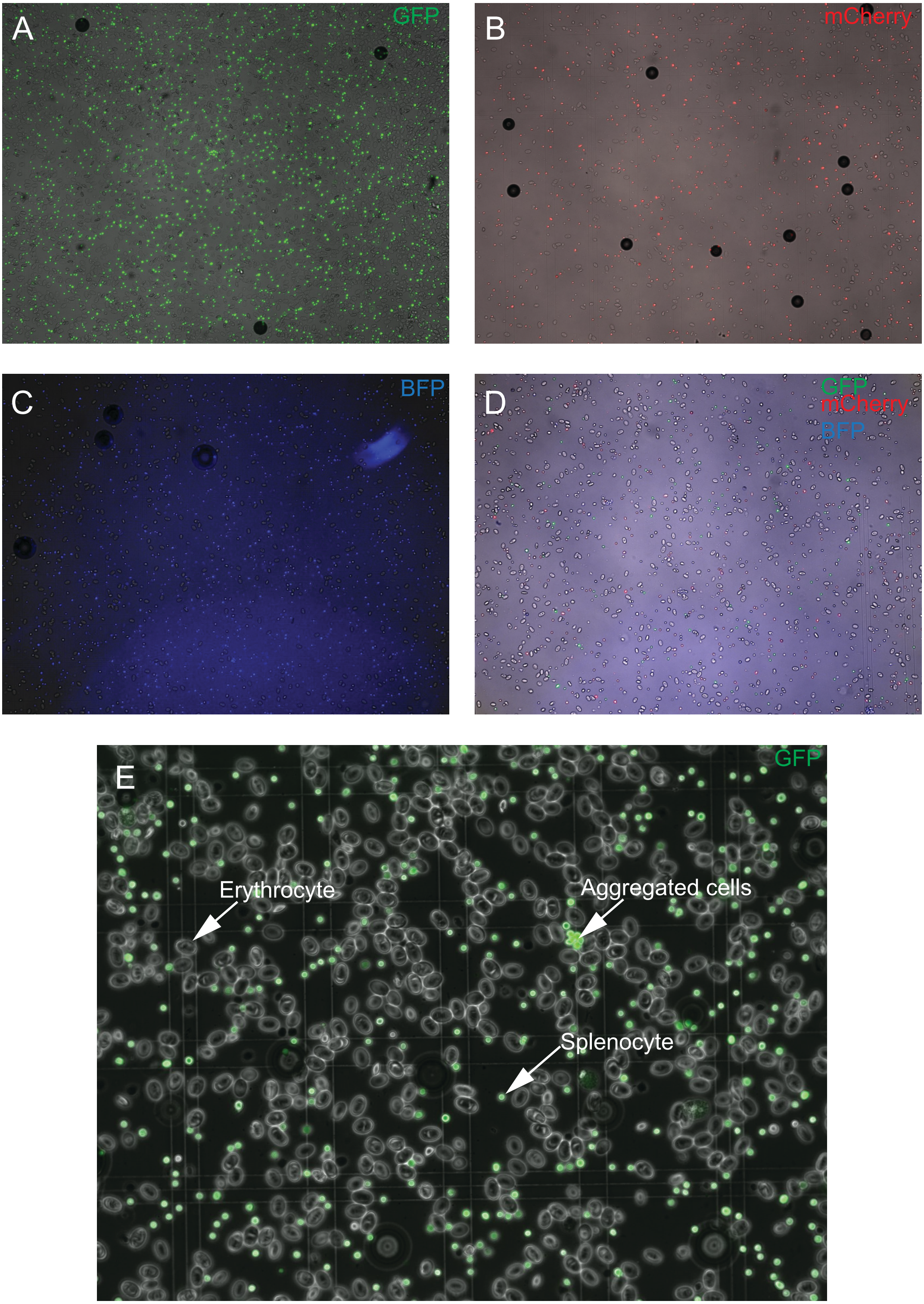


**Supplemental Figure 5:** Images of fluorescent expression of cells from fluorescent pooled scRNA-seq. Expression of GFP (A), mCherry (B), BFP (C), and a pool from these three animals (D) are depicted. (E) Splenocytes (small, round, GFP+) show robust GFP expression while erythrocytes (oval, large, GFP-) are GFP negative.

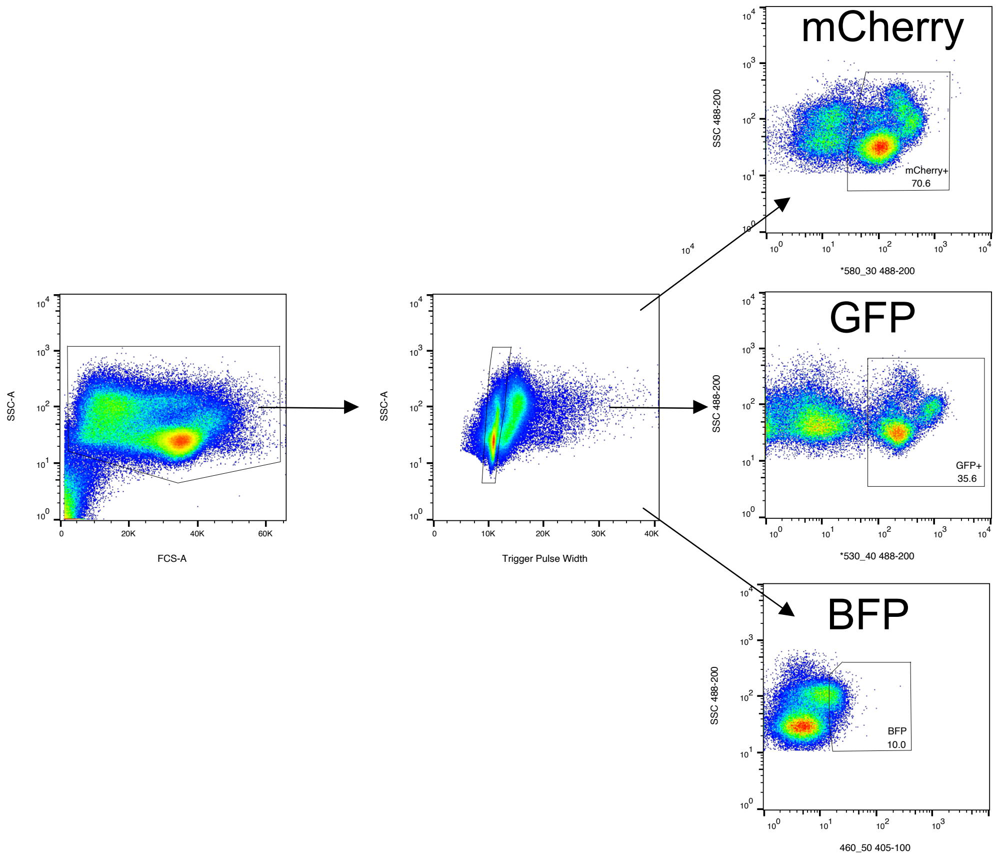


**Supplemental Figure 6:** Hierarchical gating strategy for splenocyte sorting. Cells were defined in forward-side scatter plot. Singlets as determined by location in the SSC-A vs Trigger pulse width plot were selected while removing small debris, doublets and the larger erythrocytes as defined by side scatter trace. Singlets were then plotted against relevant fluorescent markers for each transgenic animal. Labels are from animals used in the *Pleurodeles* fluorescent pooling experiment. BFP was dim but detected based on the sequencing.

A

| Dataset: | Mapping reference type used: | Common SNP file used? | Common SNP file source: | souporcell k value used: | Minimap2/freebayes run separate from souporcell pipeline? | SNP #s in VCF: |
|---|---|---|---|---|---|---|
| **Zebrafish** | genome | Yes | https://research.nhgri.nih.gov/manuscripts/Burgess/zebrafish/downloads/NHGRI-1/danRer11/danRer11Tracks/NHGRI1.danRer11.variant.vcf.gz | 3 | No | 13745235 |
| **Axolotl** | genome | Yes | http://ambystoma.uky.edu/hubExamples/hubAssembly/hub_AmexG_v6/AmexG_v6.hub_data/SNP_vcf_tracks/ddMale_to_AmexGv6.vcf.gz | 3 | No | 27191 |
| **Xenopus** | genome | No | | 8 | No | 108101 |
| **Pleurodeles** | supertranscriptome | No | | 3 | Yes | 731811 |
| **Dual salamander species** | supertranscriptome | No | | 4 | Yes | 461769 |
| **Green Monkey** | genome | Yes | https://ftp.ebi.ac.uk/pub/databases/eva/PRJEB7923/Vervet500KFinal_EVA.vcf.gz | 5 | No | 497164 |

B

| Dataset: | souporcell scripts: |
|---|---|
| Zebrafish | singularity exec Demuxafy.sif souporcell_pipeline.py -i $BAM -b $BARCODES -f $FASTA -t 20 -o $SOUPORCELL_OUTDIR -k 3 --common_variants $VCF |
| Axolotl | singularity exec Demuxafy.sif souporcell_pipeline.py -i ${BAM} -b ${BARCODES} -f ${FASTA} -t 20 -o ${OutputName} -k 3 --skip_remap SKIP_REMAP --common_variants ${VCF} |
| Xenopus | singularity exec souporcell_latest.sif souporcell_pipeline.py -i $BAM -b $BARCODES -f $FASTA -t 20 -o $SOUPORCELL_OUTDIR -k 8 |
| Pleurodeles / Dual salamander species | export SINGULARITY_BIND="${DemuxSoupDir},${MappingAnalysisDir},${FastaDir}"<br>singularity exec ${DemuxSoupDir}Demuxafy.sif renamer.py --bam $BAM --barcodes $BARCODES --out ${OutputName}.fq<br>singularity exec ${DemuxSoupDir}Demuxafy.sif minimap2 -ax splice -I 9G -t 20 -G50k -k 11 -K 50M -w 15 --sr -A2 -B8 -O12,32 -E2,1 -r200 -p.5 -N20 -f1000,5000 -n2 -m20 -s40 -g2000 -2K50m --secondary=no ${FASTA} ${OutputName}.fq > minimap.sam<br>singularity exec ${DemuxSoupDir}Demuxafy.sif retag.py --sam minimap.sam --out minimap_tagged.bam<br>singularity exec ${DemuxSoupDir}Demuxafy.sif samtools sort minimap_tagged.bam > minimap_tagged_sorted.bam<br>singularity exec ${DemuxSoupDir}Demuxafy.sif samtools index minimap_tagged_sorted.bam<br>#freebayes run:<br>singularity exec ${DemuxSoupDir}Demuxafy.sif freebayes -f ${FASTA} -iXu -C 2 -q 20 -n 3 -E 1 -m 30 --min-coverage 6 minimap_tagged_sorted.bam > free.vcf<br>#souporcell run.<br>VCF=${CurrentAnalysisDir}free.vcf<br>singularity exec ${DemuxSoupDir}Demuxafy.sif souporcell_pipeline.py -i ${CurrentAnalysisDir}minimap_tagged_sorted.bam -b ${BARCODES} -f ${FASTA} -t 20 -o ${OutputName} -k $N --skip_remap SKIP_REMAP --common_variants ${VCF} |

**Supplemental Table 1:** Bioinformatic details for souporcell demultiplexing of pooled single cell datasets. A) Table of genomic resources used as input for souporcell demultiplexing of each experiment. B) Details on souporcell scripts or computational work-through for souporcell demultiplexing of each sample. For analyses of Pleurodeles and dual salamander species datasets, the default souporcell pipeline failed for us. To address this, the first two steps of the souporcell pipeline were run separately and then the output from these was introduced back into the souporcell pipeline for completion (bottom of Table 1B). This allowed us to adjust computational parameters in the remapping stage that permitted the function to finish.
